# Tissue-specific MicroRNA Expression Alters Cancer Susceptibility Conferred by a *TP53* Noncoding Variant

**DOI:** 10.1101/582478

**Authors:** Qipan Deng, Hui Hu, Xinfang Yu, Shuanglin Liu, Lei Wang, Weiqun Chen, Chi Zhang, Zhaoyang Zeng, Ya Cao, Ling Li, Mingzhi Zhang, Steven Rosenfeld, Shideng Bao, Eric Hsi, Ken H Young, Zhongxin Lu, Yong Li

## Abstract

Patients carrying *TP53* germline mutations develop Li-Fraumeni syndrome (LFS), a rare genetic disorder with high risk of several cancers, most notably breast cancer, sarcoma, and brain tumors. A noncoding polymorphism (rs78378222) in *TP53*, carried by scores of millions of people, was associated with moderate risk of brain, colon, and prostate tumors, and other neoplasms. We found a positive association between this variant and soft tissue sarcoma (odds ratio [OR] = 4.55, *P* = 3.3 x 10^−5^). In sharp contrast, this variant was protective against breast cancer (OR = 0.573, *P* = 0.0078). We generated a mouse line carrying this variant and found that this variant accelerated spontaneous tumorigenesis and tumor development at the brain, prostate, colon, and skeletal muscle, but strikingly, it significantly delayed mammary tumorigenesis. The variant created a miR-382 targeting site and compromised a miR-325 targeting site. Their differential expression resulted in p53 downregulation in the brain and several other tissues, but p53 upregulation in the mammary gland of the mutant mice compared to that of wild-type littermates. Thus, this *TP53* variant is at odds with LFS mutants in breast cancer predisposition yet consistent with LFS mutants in susceptibility to soft tissue sarcoma and glioma. Our findings elucidate an underlying mechanism of cancer susceptibility that is conferred by genetic variation and yet altered by microRNA expression.

## Introduction

Li-Fraumeni Syndrome (LFS; also called sarcoma, breast, leukemia and adrenal gland [SBLA] syndrome in the past) is a rare, inherited familial predisposition to a wide range of cancers ^1–4^. Mutations in the coding sequence (CDS) of *TP53*, encoding the p53 tumor suppressor, are found in ~75% of LFS families; these *TP53* mutants produce mutant p53 proteins that lack most or all tumor-suppressive functions and often confer oncogenic properties ^5–7^. Changes in *TP53* noncoding sequences, in contrast, have lower penetrance but still confer cancer susceptibility. A noncoding single nucleotide polymorphism (SNP, rs78378222) in *TP53* is associated with moderate risk of several cancers^8^. Located in the fifth nucleotide of the *TP53* polyadenylation signal (PAS), the minor allele of this SNP is “*C*,” resulting in an alternative PAS (AATA***<u>C</u>***A) instead of the canonical PAS (AATA***<u>A</u>***A). Unlike LFS mutant and common *TP53* CDS variants such as P72R and P47S ^9, 10^, this *TP53* noncoding variant produces native p53 proteins, *albeit* at a lower level in cells^11^. Cancer susceptibility conferred by this *TP53* noncoding variant^8, 12–15^ does not strictly mirror that of *TP53* germline coding mutations in LFS patients^16^ (**Table S1**): individuals with the minor allele are at increased risk of brain tumors^8^ (particularly glioma^8, 12–14, 17^), neuroblastoma^18^, skin basal cell carcinoma (BCC)^8^, esophageal squamous cell carcinoma (SCC) ^19^, prostate cancer, colorectal adenoma^8^, and uterine leiomyoma^20^.

In LFS patients, the risk of developing any invasive cancer (excluding skin cancer) is approximately 50% by age 30 (compared with 1% in the general population), and approximately 90% by age 70 ^21^. Although numerous tumor types are seen in patients with LFS, five core cancers (breast cancer, soft-tissue sarcoma, osteosarcoma, brain tumor, and adrenocortical carcinoma) make up about 80% of LFS-associated tumors^1–3^. Brain tumors occur in 9-16% of LFS patients, with glioma being the most common (>40%)^22, 23^. Sporadically, glioma accounts for ~80% of all primary adult malignant brain tumors. That the risk of glioma is increased 2-fold in relatives of glioma patients provides evidence for an inherited risk ^24^. A number of rare inherited cancer predisposition disorders such as LFS, Turcot syndrome, and neurofibromatosis are recognized to be associated with increased risk of glioma. Scores of common SNPs were recently identified as increasing the risk of glioma^15, 17, 25^. Supported by 5 independent studies^8, 12–15, 17^, the *TP53* noncoding variant (rs78378222[C]) increases glioma risk more significantly than for other tumors with an odds ratio (OR) ranging from 2.35 to 3.74 (**Table S1**). It is estimated that this variant alone could represent up to 6% of the familial risk of glioma^13^. Thus, this *TP53* variant shares a phenotypic similarity with LFS mutants in brain tumor (glioma) predisposition.

Breast cancer is the number one tumor type found in adult LFS patients. Nearly 80% of LFS females develop breast cancer, whereas no LFS male does^22^. Even when male and female patients are considered together, breast cancer is found in 39% of patients, while soft tissue sarcoma (the second most common tumor type) is found in 27% of patients^22^. This *TP53* noncoding variant does not appear to increase the risk for sporadic breast cancer, nor for high-risk breast cancer (**Table S1**)^8^; however, in this study, all breast cancer patients and the vast majority of the unaffected controls were genotyped for this variant by imputation^8^, which has relatively low accuracy for infrequent alleles like this *TP53* variant^26^. In fact, for a variant with a frequency < 2%, there is no such imputation method that achieves ≥ 95% concordance with Taqman real-time PCR or DNA sequencing^26–28^. In addition, other cancers to which patients with this variant are predisposed (neuroblastoma, prostate cancer, skin BCC, and esophageal SCC) occur infrequently in patients with LFS ^3, 29, 30^.

In this study, we performed direct genotyping of this *TP53* variant in patients with breast cancer and sarcoma. We uncovered that this *TP53* variant increased the risk for soft tissue sarcoma, but decreased the risk for breast cancer. We then generated a mouse line carrying this variant and evaluated tumorigenesis in different organs; particularly, we investigated whether and how this variant increases the risk for glioma but not breast cancer. We found that this *TP53* variant created a targeting site for miR-382-5p (miR-382) that was highly expressed in the brain and compromised the site for miR-325-3p (miR-325) that was highly expressed in the mammary gland. Differential expression of these 2 microRNAs (miRNAs) was likely responsible for observed p53 upregulation in the mammary gland, but p53 downregulation in the brain and several other tissues in mutant mice as compared to wild-type littermates. Our findings uncover a *TP53* variant at odds with LFS mutants in regard to breast cancer risk yet consistent with LFS mutants in predisposition to soft tissue sarcoma and glioma and reveal an underlying mechanism of tissue-specific cancer susceptibility that is mediated by miRNAs.

## Results

### Susceptibility to breast cancer and sarcoma

To determine the association of this *TP53* noncoding variant with the risk for breast cancer and sarcoma, we genotyped rs78378222 in adult patients with breast cancer (n=2,373) or sarcoma (n=130) and their respective unaffected controls (n=9,972 and 8,947) of Chinese Han descent (**Table 1**). The minor allele frequencies (MAFs) in the controls were 0.017-0.018, comparable to previous reports (**Table S1**). We observced a significant association between rs78378222[C] and the risk of sarcoma (OR = 3.29, *P* = 0.0014). The risk was limited to soft tissue sarcoma (OR = 4.55, *P* = 3.3 x 10^−5^) because no patients with osteosarcoma carried this variant. In sharp contrast, this *TP53* variant protected against breast cancer (OR = 0.573, *P* = 0.0078).

**Table 1.** Association between breast cancer and sarcoma and rs78378222[C].

| Cancer Types | OR | *P* | 95% CI | Cases |  | Controls |  |
| --- | --- | --- | --- | --- | --- | --- | --- |
|  |  |  |  | N | Frequency | N | Frequency |
| Breast cancer | 0.573 | 0.0078 | 0.374-0.867 | 2,373 | 0.0107 | 9,972 | 0.0180 |
| Sarcoma | 3.29 | 0.0014 | 1.51-7.17 | 130 | 0.0538 | 8,947 | 0.0170 |
| Soft tissue sarcoma | 4.55 | $3.3 \times 10^{-5}$ | 2.07-9.99 | 96 | 0.0729 | 8,947 | 0.0170 |
| Osteosarcoma | NA | NA | NA | 34 | 0 | NA | NA |

OR, odds ratio; CI, confidence interval; NA, not applicable; Controls were patients who were not diagnosed with any cancer.

### A mouse model for this variant and spontaneous tumorigenesis

To investigate the causative nature of this variant in cancer, we generated a mouse line with a *Trp53* gene carrying the alternative PAS (**Fig. S1a,b**). We termed the resultant mutant *Trp53* allele *Trp53*^1755C^ (abbreviated as *p53*^C^) because the 1755^th^ nucleotide of mouse *p53* reference mRNA is the ortholog of human rs78378222. Mice carrying the *p53*^C^ allele had shorter survival than their WT *p53*^+/+^ littermates (**Fig. 1a**), with median survival of 108 weeks for heterozygous (*p53*^+/C^) and 102 weeks for homozygous (*p53*^C/C^) mice, but longer survival than *p53*^+/−^ mice (a median survival of ~70 weeks)^31^. Tumors developed more frequently in mutant mice than in WT controls (**Fig. S1c**). B-cell malignancies were the most frequently occurring cancer in mutant mice (without development of thymic lymphoma) (**Fig. 1b-1e** and **Table S2**). Exome sequencing of lymphomas in the mutant mice showed *p53* coding missense and truncation mutations in all tumors, with a relative read ratio of *p53* and *Gapdh* close to 1:1 (**Table S3**, **Fig. S1d**). Irradiation further shortened the survival of *p53* mutant mice, with increased cancer-associated death and increased thymic lymphoma incidence compared with untreated mice (**Fig. S1e-h**). These data indicate that this p53 noncoding variant increases the risk of malignancies and shortens survival.

**Fig. 1.** The *p53^C^* allele accelerates spontaneous tumorigenesis and *Pdgfb*-driven gliomagenesis in mice. (**a**) Survival of wild type (WT, *p53*^+/+^), heterozygous (*p53*^+/C^), and homozygous (*p53*^C/C^) littermate mice. *P* < 0.0001 (log-rank test). (**b**) Hematoxylin and eosin stain (H&E) staining of representative follicular lymphoma (FL) and splenic marginal zone lymphoma (SMZL) in mutant mice. Scale bar, 200 μm (top row), 20 μm (bottom row). (**c**) Diffuse large B-cell lymphoma (DLBCL, top row) and histiosarcoma (bottom row) in mutant mice. Scale bar, 20 μm. (**d**) Immunohistochemistry (IHC) staining for B cell marker B220 in FL and SMZL in mutant mice. Scale bar, 200 μm (top row), 20 μm (bottom row). (**e**) IHC staining for plasma cell marker Ig κ light chain of SMZL in mutant mice. Scale bar, 200 μm (top row), 50 μm (bottom row). (**f**) Survival of mice with different *p53* alleles injected in the right subventricular zone (SVZ) with retrovirus carrying *Pdgfb*. *P* < 0.0001 (log-rank test). (**g**) H&E staining of glioma in *p53*^C/C^ mice (*n*=9). **1**, a representative tumor shown at low power magnification (2X). Scale bar, 1 mm. **2**, invasive growth of the tumor cells. Arrows showing invasion front of the tumor. Scale bar, 100 μm. **3**, high power view (1000X) of a blood vessel (arrows) in the tumor tissue. **4**, infiltrating monocytes (arrow) within the tumor. **5**, necrosis in the tumor tissue. Arrows point to necrotic cells. **6**, highly invasive tumor cell (arrows) growth surrounding a neuron (star). Scale bar in 3-6, 5 μm. (**h)** IHC staining for the astrocyte/glial cell marker GFAP. Arrow showing a tumor cell staining positive for GFAP (dark brown, right), but most tumor cells lost GFAP expression. (**i**) IHC staining of the neuronal marker synaptophysin. Scale bar in (**h** and **i**), 100 μm (left), 10 μm (right). (**j**) IHC analyses of glioma stem cell markers in *p53*^C/C^ mice. *Pdgfb* was tagged with 3x hemagglutinin (HA). Staining of HA for *Pdgfb* expression, PCNA for tumor cell proliferation, neuralgia progenitor/glioma stem cell makers GD3 (red arrows point to the accumulation of GD3 at invasive front line; blue arrow points to necrotic cells at the GD3-positive invasive front line), Nestin, and Oligo2. Scale bars: 1 mm (top row), 100 μm (second row), and 5 μm (bottom row). (**k**) IHC analyses of other glioma markers in *p53*^C/C^ mice. IHC staining was performed for the astrocyte/glial cell marker GFAP, p53, and the epithelial-mesenchymal transition (EMT) markers β-catenin, vimentin, and α-SMA in *Pdgfb*- induced glioma obtained from *p53*^C/C^ mice. In the bottom row, the green arrow indicates a GFAP-positive cell with typical astrocyte morphology, the red arrow indicates a tumor cell with strong GFAP-positive staining. Scale bars: 100 μm (top row), 20 μm (second row), and 5 μm (bottom row). N, normal tissues; T, tumors; images were representative of staining obtained in 6 tumors (1 per animal) (**g-k**). (**l**) Survival of syngeneic WT mice orthotopically transplanted with glioma cells (1, 3, or 6 x 10^5^) isolated from tumors developed in *p53*^C/C^ mice (**f**). *P* = 0.003 (log-rank test).

### Glioma

*TP53* is the only gene that 1) causes a rare monogenic Mendelian disorder (LFS) that includes a greatly increased risk of glioma, and 2) has a common inherited SNP (rs78378222) associated with a smaller increased risk of glioma (**Table S1**). Glioblastoma (GBM) is the highest grade astrocytoma. This *TP53* variant predispose both GBM and non-GBM glioma^17^. Approximately 30% of GBM tumors demonstrate a proneural gene expression pattern, characterized by frequent *TP53* mutation and *PDGFRA* mutation and/or overexpression ^32^. We employed a PDGF-driven GBM mouse model with a proneural gene expression pattern by injecting a retrovirus expressing *Pdgfb* into the brain subventricular zone (SVZ) ^33^. *p53*^−/−^ mice receiving retrovirus had the earliest tumor onset, and all succumbed to glioma within 40 days, whereas *p53*^+/−^ mice survived up to 150 days (**Fig. 1f**). All *p53*^C/C^ mice succumbed to glioma between 45 and 260 days, but neither *p53*^+/C^ nor *p53*^+/+^ mice with viral *Pdgfb* had developed any tumors by day 275 when they were all sacrificed, and none showed any brain lesions. Tumors from *p53*^C/C^ mice were highly infiltrative and invasive with extensive vascularization and necrosis (**Fig. 1g**), all histological hallmarks of GBM. These tumors extensively expressed markers for GBM stem cells and the epithelial-mesenchymal transition (EMT) (**Fig. 1h-1k**). The malignant phenotype of tumor cells was further demonstrated by transplanting them into syngeneic immunocompetent recipient mice in a dose-dependent manner (**Fig. 1l**). These data support that the *TP53* variant is pathogenic in gliomagenesis.

### Soft tissue sarcoma

We also determined the role of this *p53*^C^ allele in regulating p53 in mouse embryonic fibroblast cells (MEFs) and in chemically induced sarcomagenesis in mice as this *TP53* variant significantly increase the risk for soft tissue sarcoma (**Table 1**). *p53*^C^ reduced p53 expression and activities in MEFs as judged by multiple molecular and cellular assays, but it did not alter the half-life of p53 protein (**Fig. S2**). When exposed to 3-methylcholanthrene, *p53*^C/C^ and *p53*^+/C^ mice developed tumors faster and had shorter survival than *p53*^+/+^ mice (**Fig. S3** and **Table S4**). Together, these data demonstrate that *p53*^C^ reduces p53 expression and function in MEFs and promotes chemical-induced sarcomagenesis.

### Breast cancer

To determine the role of this *TP53* variant in the pathogenesis of breast cancer, we first crossed *p53*^+/C^ mice with mice expressing the polyoma virus middle T antigen (*PyVT*) driven by the mouse mammary tumor virus long terminal repeat (MMTV) promoter in the mammary gland ^34^. To our surprise, *p53*^+/C^;*PyVT* F1 females developed smaller and fewer mammary tumors and survived longer than *p53*^+/+^;*PyVT* littermates (**Fig. 2a-c**). We examined the precancerous transformation of mammary epithelial cells using whole mounts of mammary gland and found that the *p53*^C^ allele delays development of epithelial hyperplasia to about 8 weeks of age, compared to 4 to 6 weeks in *p53*^+/+^;*PyVT* littermates (**Fig. 2d**). In F2 offspring, both *p53*^C/C^;*PyVT* and *p53*^+/C^;*PyVT* mice had longer tumor-free survival than *p53*^+/+^;*PyVT* littermates (**Fig. 2e**). Tumors from mutant mice were more differentiated and showed less EMT compared to tumors from *p53*^+/+^;*PyVT* mice at the same age, as evidenced by the higher expression of β-casein and E-cadherin (**Fig. S4a**).

**Fig. 2.**
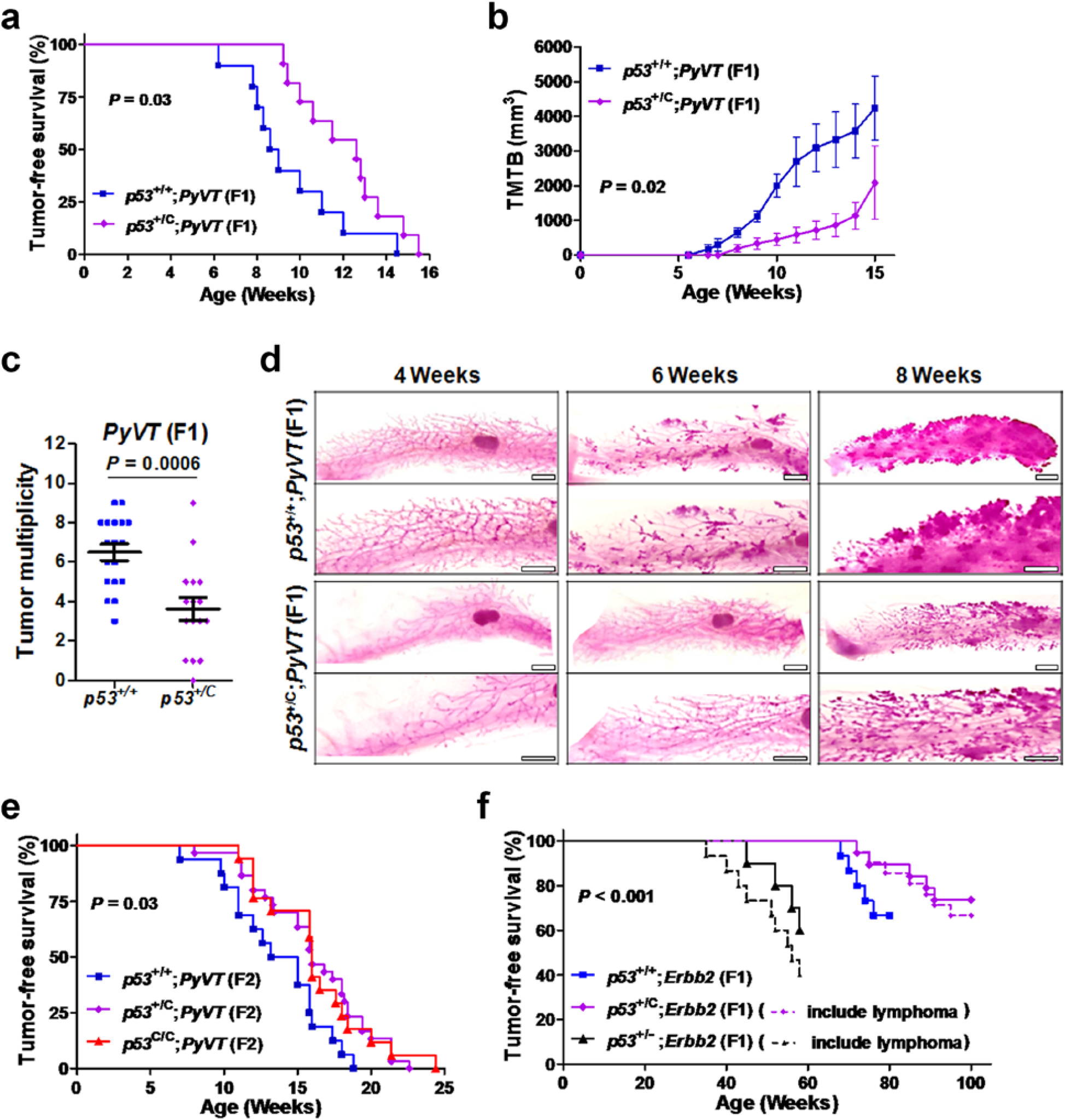
The *p53*^C^ allele delays either *PyVT*- or *Errb2*-driven mammary tumorigenesis in mice. (**a**) Tumor-free survival of F1 female offspring of *p53*^+/C^ mice crossed with *PyVT*- overexpressing mice (FVB/N-Tg(MMTV-PyVT)634Mul/J). *n*=10 for *p53*^+/+^;*PyVT* and n=11 for *p53*^+/C^;*PyVT*; *P* = 0.03 (log-rank test). (**b**) Tumor volume in F1 female *p53*^+/+^;*PyVT* and *p53*^+/C^;*PyVT* mice at indicated ages. TMTB, total measured tumor burden. *P* = 0.02. (**c**) Tumor multiplicity in F1 females at 15 weeks. Data were presented as mean ± s.e.m; *P* = 0.0006 (Mann-Whitney *U* test). (**d**) Whole-mount carmine red stain of the mammary glands from F1 females at indicated ages. *n* = 5 mice per group. Scale bar, 2 mm. (**e**) Tumor-free survival of F2 female offspring (obtained by crossing F1 *p53*^+/C^;*PyVT* with F1 *p53*^+/C^ mice). *P* = 0.03. (**f**) Mammary tumor-free survival and tumor-free survival of F1 female offspring from *Erbb2* mice interbred with *p53*^+/C^ mice. As *p53*^+/C^ and p53^+/−^ mice may develop lymphoma, dashed lines indicated tumor-free survival (*P* = 0.006) and solid lines indicated mammary tumor-free survival (*P* < 0.001), i.e., dashed lines included 2 of the 21 *p53*^+/C^;*Erbb2* mice and 5 of the 15 p53^+/−^;*Erbb2* mice that developed lymphomas. *p53*^+/−^;*Erbb2* mice, as a control, were obtained by breeding *p53*^−/−^ mice with *Erbb2* mice.

As most breast tumors that develop in LFS females are ERBB2-positive^22, 35^, we next crossed *p53*^+/C^ mice with mice overexpressing MMTV-driven *Erbb2*. Similarly, *p53*^+/C^;*Erbb2* mice had longer tumor-free survival than the *p53*^+/+^;*Erbb2* littermates, whereas *p53*^+/−^;*Erbb2* mice had the shortest tumor-free survival (**Fig. 2f**). Tumors from *p53*^+/C^;*Erbb2* mice expressed higher levels of E-cadherin and pan cytokeratin (PCK) than the *p53*^+/+^;*Erbb2* control, indicating their epithelial origin and slower cancer progression in *p53* mutant mice (**Fig. S4b,c**). *p53*^+/C^;*Erbb2* mice displayed less severe lung metastasis with smaller metastasized tumors, lower incidence of metastasis, and less multiplicity than *p53*^+/+^;*Erbb2* littermates (**Fig. S4d,e,f**). p53 protein staining was strong in mammary tumors that developed in mice with either the mutant or WT *p53* (**Fig. S5**). DNA sequencing of the *p53* gene in 5 mammary tumors from each group revealed all samples carried heterozygous *p53* coding mutations, supporting that gain-of-function p53 mutation is needed for mammary tumorigenesis.

### p53 deregulation and differential miRNA expression

We analyzed p53 expression and activation in various tissues. p53 protein and mRNA levels, as well as protein phosphorylation upon irradiation, in brain, colon, and spleen of *p53*^C/C^ mice were lower than in WT littermates (**Fig. 3a-c**). p53 protein in the brain and skeletal muscle of *p53*^C/C^ mice was significantly lower than that in *p53*^+/+^ mice (**Fig. 3a**), in line with observed gliomagenesis (**Fig. 1f**) and accelerated sarcomagenesis in *p53*^C/C^ mice (**Fig. S3**). In contrast, p53 was upregulated in the mammary gland of *p53*^C/C^ mice compared to WT littermates. When cDNA was prepared from total RNAs in mouse tissues and sequenced, the *p53*^C^ alleles from *p53*^C/C^ mice were outnumbered by *p53*^A^ alleles from *p53*^+/+^ mice in the brain, whereas the *p53*^C^ allele outnumbered *p53*^A^ in the mammary gland (**Fig. 3d**). We cloned the whole native human *TP53* 3′UTR (the *A* allele, *TP53*^A^) or the 3′UTR with the alternative PAS (the *C* allele, *TP53*^C^) downstream to a EGFP or RFP (*EGFP*^C^ and *RFP*^A^) driven by the *TP53* promoter. These two constructs were introduced into cells and the mean fluorescence intensity (MFI) and EGFP/RFP mRNA levels were determined. Both normalized MFI and mRNA of *EGFP*^C^ were lower than that of *RFP*^A^ in U87 glioma cells, yet the opposite was observed in MCF10a and breast cancer MDA-MB-231 cells (**Fig. 3e, f**). When *EGFP/RFP* was replaced by the *TP53* coding sequence, the *A* allele resulted higher p53 expression than the *C* allele in U87, but not in MCF7 cells (**Fig. 3g**). Thus, the variant resulted in p53 upregulation or downregulation, dependent on the tissue or cellular context.

**Fig. 3.**
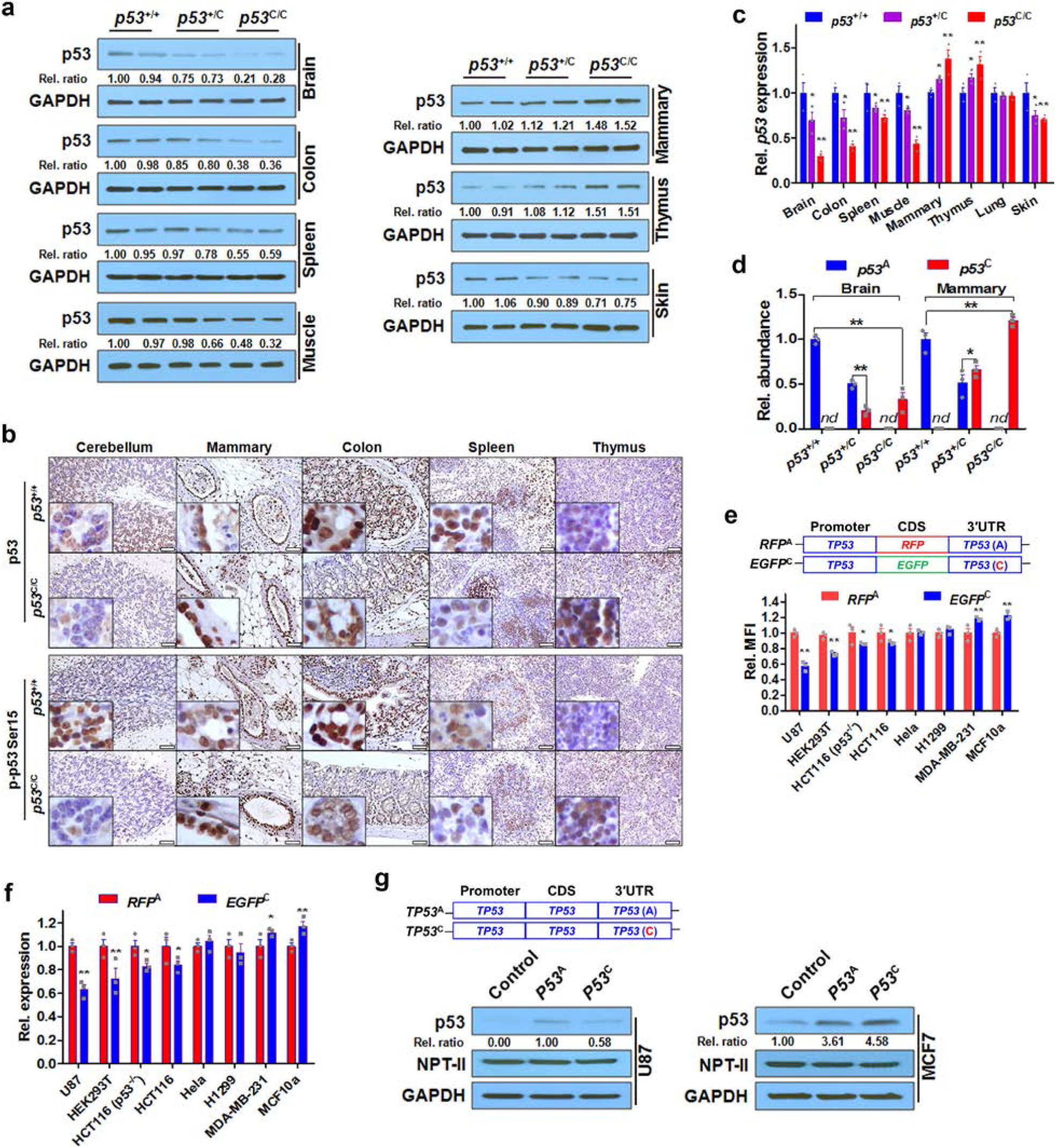
Differential p53 deregulation by the *p53*^C^ allele in tissues and cell lines. (**a**) Western blotting analyses for p53 protein levels in tissues from mouse littermates. Each pair of lanes indicates samples from two different mice. Rel. ratio: the relative signal densities of p53 as referenced to that of GAPDH. (**b**) IHC staining for p53 (upper panel) and phosphorylated p53 at site Ser15 (lower panel) in *p53*^+/+^ and *p53*^C/C^ mice. Mice were treated with 4 Gy total body γ-irradiation and brain, mammary gland, colon, spleen and thymus were collected 6 h later for IHC staining. Scale bar, 50 μm. *n* = 3 mice per group, inserts show high-magnification (400X) images. (**c**) Quantitative real-time PCR (qPCR) analysis for *p53* mRNA levels in mouse tissues. Rel. *p53* expression: Relative *p53* mRNA levels normalized to *Gapdh*. Data were presented as mean ± s.e.m; *n* = 3 independent biological replications. (**d**) Relative abundance (Rel. abundance) of “A” and “C” p53 cDNA alleles as determined by sequencing the PCR product of p53 cDNAs reverse transcribed from total mRNAs extracted from the brain and mammary tissues of mouse littermates. *nd*, not detectable. (**e**) The relative mean fluorescence intensity (Rel. MFI) of *EGFP*^C^ *vs RFP*^A^ in several cell lines transfected with 2 constructs as determined by FACS. CDS, coding sequence. Data were presented as mean ± s.e.m; *n* ≥ 3. (**f**) qPCR analysis for *EGFP/RFP* mRNA levels from indicated cell lines transfected with *EGFP*^C^ or *RFP*^A^, normalized to that of *neomycin phosphotransferase* (NPT-II), a gene carried by the vector. Rel. expression: Relative expression. Data were presented as mean ± s.e.m; *n* = 3 biological triplicates for each cell line. (**c-f**) *, *P* < 0.05, **, *P* < 0.01 (one-way ANOVA). (**g**) Western blotting analyses for p53 protein levels in MCF7 and U87 cells transfected with a plasmid expressing *TP53* with a WT 3′UTR (*TP53*^A^) or a 3′UTR with an alternative PAS (*TP53*^C^). Rel. ratio: the relative signal densities of p53 as referenced to that of NPT-II. *n* = 3 biological triplicates for each cell line.

Because the variant exerts moderate regulation over p53, we suspected that miRNA expression is a determinant of differential p53 deregulation in mutant mice. miR-325 putatively targets the native *TP53* 3′UTR, whereas miR-382 targets the *TP53* 3′UTR with the PAS variant in both human and mouse (**Fig. 4a** and **Fig. S6a**). miR-325 and miR-382 expression varies markedly across different human tissues with miR-325 highly expressed in the breast and miR-382 highly expressed in the brain (**Fig. S6b-f**). The ratio of miR-325: miR-382 is lowest in the brain and highest in the breast/mammary gland in both humans (**Fig. 4b**) and mice (**Fig. 4c**). Like miR-325, three other miRNAs putatively targeting *TP53*^A^ are also highly expressed in human breast tissues compared to brain (**Fig. S7**). The *p53*^C^ allele itself did not alter the expression of miR-325 or miR-382 in the brain, mammary gland, and other tissues in mice (**Fig. S6g,h**). We next examined the expression of these 2 miRNAs in several human cell lines: U87 glioma and HCT116 colon cancer cells had the lowest miR-325: miR-382 ratio, compared to MCF10a, MCF7, and two other breast cancer cell lines (**Fig. 4d**, **Fig. S6i, j**). We then determined miRNA regulation of endogenous p53 in these cells. miR-382 inhibition resulted in higher p53 protein expression than did miR-325 inhibition in U87 and HCT116 cells, which had higher endogenous miR-382 levels, while opposite results were observed in MCF7 and MCF10a cells, which had higher endogenous miR-325 levels (**Fig. 4e**). Similarly, miR-325 inhibition resulted in greater p53 protein expression than did miR-382 inhibition in *p53*^+/+^ MEFs, while opposite results were observed in *p53*^C/C^ MEFs (**Fig. 4f**).

**Fig. 4.**
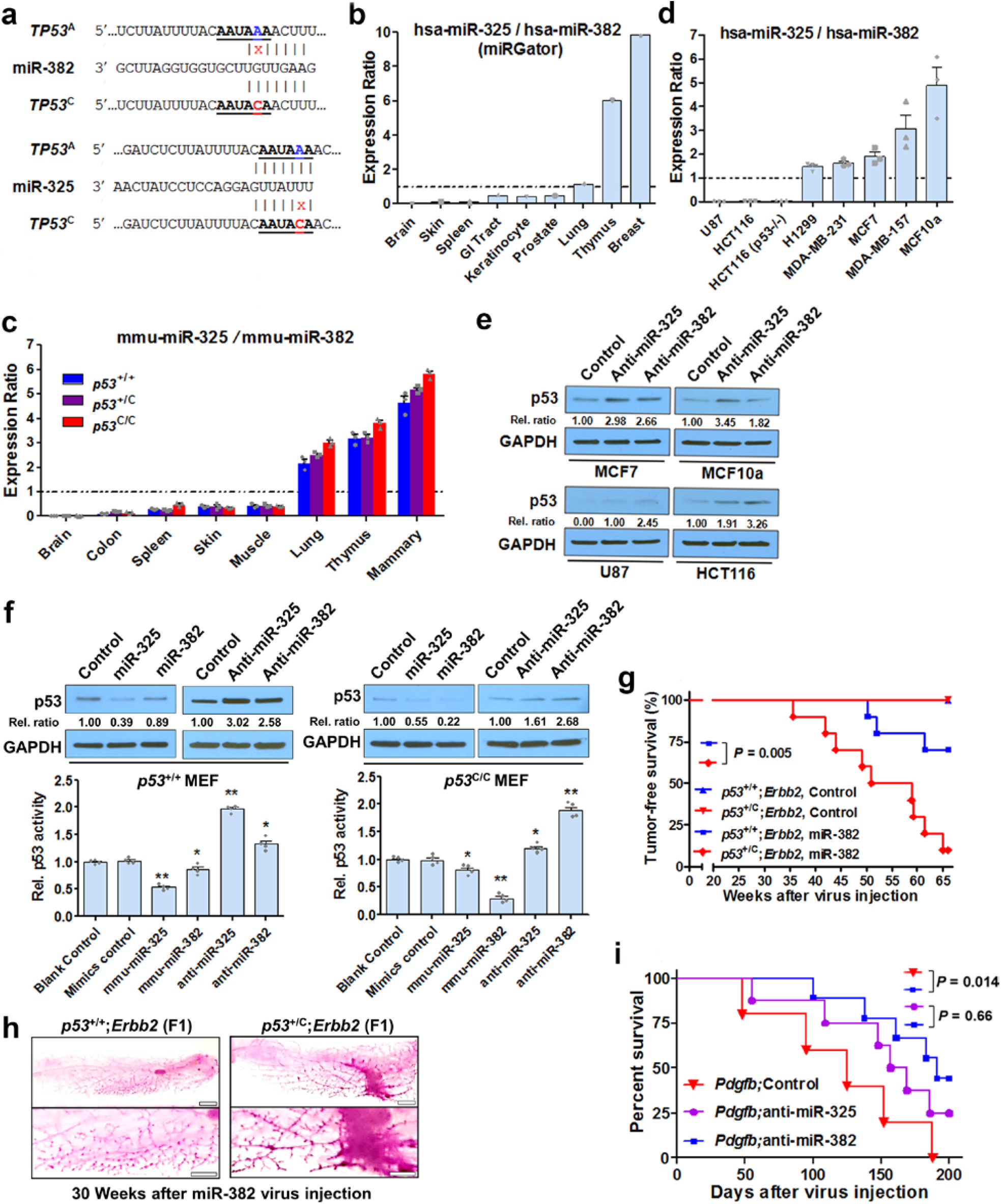
Tissue-specific miRNA expression results in differential *p53* deregulation in the brain and mammary tissues from mice with the *p53*^C^ allele. (a) Schematic diagram showing that human *TP53* variant (in red, *TP53*^C^) creates a miR-382 (hsa-miR-382-5p) targeting site and compromises the site on native *TP53* (*TP53*^A^) targeted by miR-325 (hsa-miR-325-3p). miRNA targeting was predicted by both TargetScan and PolymiRTS. (**b**) miR-325: miR-382 expression ratio in different human tissues, calculated by dividing the mean miR-325 read count by the mean miR-382 read count, using deep sequencing data obtained from the online database miRGator. (**c**) miR-325: miR-382 expression ratio in different mouse tissues using qPCR. (**d**) miR-325: miR-382 expression ratio in the indicated human cancer cell lines using qPCR. Data in (**c,d**) were presented as mean ± s.e.m; *n* = 3 biological triplicates for each cell line with 3 technical triplicates. (**e**) Representative Western blotting analysis for p53 expression in human cell lines. (**f**) p53 expression (top) and reporter activity (bottom) in *p53*^+/+^ and *p53*^C/C^ MEFs transfected with miR-325 or miR-382 inhibitors or mimics. In **e** and **f**, “Control” denotes the negative control #1 for mirVana miRNA inhibitors or mimics, correspondingly (ThermoFisher); Rel. ratio: the relative signal density of p53 as referenced to that of GAPDH (values were the average of 3 independent experiments). (**g**) Percent survival of *p53*^+/+^;*Erbb2* and *p53*^+/C^;*Erbb2* F1 female offspring overexpressing miR-382 in the mammary gland. Twenty-week-old mice (*n*=10 per group) were injected in the third pair of mammary glands with lentivirus carrying *miR- 382* or the control. *P* = 0.005. (**h**) Whole-mount carmine red stain of the mammary glands from *p53*^+/+^;*Erbb2* and *p53*^+/C^;*Erbb2* F1 females injected with lentivirus expressing miR-382. *n*=4 mice per group. (**i**) Survival of *p53*^C/C^ mice overexpressing *Pdgfb* and an inhibitor to miR-382. Mice were injected in the right SVZ with retrovirus expressing both *Pdgfb* and anti-miR-382 (n=9), anti-miR-325 (n=8), or an inhibitor control (n=5). *P* = 0.014 (log-rank test).

We introduced miRNA mimics into MEFs and found miR-325 downregulated p53 to a greater extent than did miR-382 in *p53*^+/+^ MEFs, whereas miR-382 downregulated p53 to a greater extent than did miR-325 in *p53*^C/C^ MEFs (**Fig. 4f**). In human cells with exogenous *TP53*^A^, miR- 325 downregulated p53 protein levels more than miR-382; in contrast, in cells with *TP53*^C^, miR- 382 downregulated p53 protein levels more than miR-325 (**Fig. S8a-d**). In ensuing reporter assays, modulation of miR-382 expression had a greater impact on *TP53*^C^-driven reporter activity than did miR-325 modulation, whereas modulation of miR-325 expression had a greater impact on *TP53*^A^-driven reporter activity than did miR-382 modulation (**Fig. S8e-m**, **Fig. 4f**). Taken together, these data suggest that miR-325 targets the *A* allele more effectively than does miR-382, whereas miR-382 targets the *C* allele more effectively than does miR-325.

To test the contribution of miRNA targeting sites (i.e., the PAS plus its adjacent nucleotides), we cloned the *TP53* coding sequence driven by the human *TP53* promoter, upstream of the SV40 3′UTR with either the WT PAS (A) or the PAS variant (C) (**Fig. S9a-b**), neither of which are targeted by miR-382. These 2 plasmids were co-transfected into five cell lines, along with a p53 activity reporter construct. Regardless of the endogenous levels of miR-382, all five cell lines with the WT PAS displayed higher p53 activity than those with the alternative PAS (**Fig. S9c**). Next, we generated EGFP-expressing plasmids with the *TP53* 3′UTR carrying either the WT PAS (A) or the PAS variant (C). We mutated the four nucleotides immediately downstream of the PAS (M4) to eliminate the miR-382 targeting site (**Fig. S9d-g**). In U87 cells with high endogenous miR-382 levels (**Fig. 4d**), the *A* allele resulted in higher EGFP expression than did the *C* allele in U87 cells. When miR-382: *TP53* 3′UTR (C) was disrupted, cells with the M4 *C* allele had higher EGFP expression than those with the M4 *A* allele (**Fig. S9h**). These results support that the integrity of the miR-382 binding site is essential to miR-382-mediated suppression of *TP53*^C^.

To further test direct interaction between *TP53*^A^/*TP53*^C^ and miR-325/miR-382, we performed pull-down assays using H1299 cells transfected with biotinylated RNA fragments and streptavidin beads to co-purify their binding partners (**Fig. S10**). First, biotinylated *TP53* 3′UTR fragments containing miRNA binding sites were used as baits to co-purify potential binding miRNAs (**Fig. S10a**). The fragment containing the native *TP53* PAS pulled down more miR-325 than miR-382, whereas the one containing the alternative PAS pulled down more miR-382 than miR-325; when the binding sites for miR-325, miR-382, or both were mutated, the respective fragment was unable to pull down miR-325, miR-382, or either of them (**Fig. S10b-d**). Second, biotinylated miRNAs were used to co-purify exogenous 3′UTR in H1299 cells transfected with EGFP-expressing plasmids containing the *TP53* 3′UTR carrying either the WT PAS or the alternative PAS with a native or mutant miR-382 targeting site (**Fig. S9d, f**; **Fig. S10e**). The 4 plasmids resulted in comparable 3′UTR levels (**Fig. S10f**). Biotinylated miR-325 pulled down the native 3′UTR more effectively than did miR-382, while miR-382 pulled down the 3′UTR with the alternative PAS more effectively than did miR-325; elimination of the miR-382 binding site completely prevented the 3′UTR from being co-purified with miR-382 (**Fig. S10g**). These experiments provide added evidence to support direct binding between *TP53*^A^/*TP53*^C^ and miR- 325/miR-382.

### *p53*^C^ suppression by miR-382 *in vivo*

We next altered miRNA expression in the mouse brain and mammary gland and evaluated changes in tumorigenesis. We chose miR-382 because it is highly abundant than miR-325 (**Fig. S6**) and has no orthologs like miR-325 (**Fig. S7a**). First, we injected a retrovirus overexpressing miR-382 into the mammary gland of F1 littermates with either *p53*^+/+^*;Erbb2* or *p53*^+/C^*;Erbb2*. By 65 weeks, miR-382 significantly accelerated mammary tumor development in *p53*^+/C^;*Erbb2* mice but had smaller impact on *p53*^+/+^;*Erbb2* mice (**Fig. 4g**). *p53*^+/+^*;Erbb2* or *p53*^+/C^*;Erbb2* mice receiving control virus did not develop tumors. With miR-382 overexpression in the mammary gland, *p53*^+/C^;*Erbb2* mice had shorter tumor-free survival than *p53*^+/+^;*Erbb2* mice (**Fig. 4g**), in marked contrast to their untreated counterparts (**Fig. 2f**). Before tumors were visible, miR-382 overexpression in the mammary gland of *p53*^+/C^;*Erbb2* mice promoted mammary epithelial outgrowth compared to that in *p53*^+/+^;*Erbb2* mice (**Fig. 4h**). Second, we injected a retrovirus overexpressing both *Pdgfb* and an inhibitor to miR-382 into the SVZ of *p53*^C/C^ mice. Those receiving both *Pdgfb* and the miR-382 inhibitor prolonged the survival of *p53*^C/C^ mice compared with those receiving *Pdgfb* and the control inhibitor (or the miR-325 inhibitor) (**Fig. 4i**). These data demonstrate that modulating the expression of *p53*^C^-targeting miR-382 alters tumor development both in the brain and in the mammary gland of *p53*^C^ mutant mice.

### Colon and prostate tumors

Lastly, we determined whether this *TP53* variant is pathogenic in colorectal adenoma and prostate cancer due to the high incidence of these malignancies worldwide. Mutant mice treated with dextran sulfate sodium developed severe colitis, as evident by massive edema, lymphocyte infiltration and loss of colonic epithelial architecture (**Fig. S11a**), and by other indicators of systemic inflammation and tissue damage (**Fig. S11b-n**). The addition of azoxymethane (AOM) induced development of colorectal adenomas that had activated β-catenin signaling in mutant mice but not in WT mice 8 weeks after treatment (**Fig. S12** and **Fig. S13a**). p53 signaling was downregulated in colon tissues from mutant mice without treatment (**Fig. S13b-c**) and these tissues exhibited higher expression of inflammatory genes (**Fig. S13d**). Next, we crossed *p53*^+/C^ mice with Hi-Myc mice carrying a prostate-specific *Myc* transgene. *p53*^C^ accelerated *Myc*-driven prostate tumorigenesis. Both Hi-Myc;*p53*^+/+^ and Hi-Myc;*p53*^+/C^ mice developed adenocarcinoma at 6 months; however, tumors from Hi-Myc;*p53*^+/C^ mice were more advanced and poorly differentiated, as suggested by cytoplasmic hyperchromasia, loss of extracellular matrix, and frequent nuclear atypia (**Fig. S14**). Collectively, these data show that *p53*^C^ promotes development of both colon and prostate tumors.

## Discussion

The architecture of inherited cancer susceptibility is a montage of predisposition alleles with different levels of risk and prevalence (i.e., allele frequency) in the general population^36^. At one end of the spectrum are very rare to rare high-penetrance mutants causing Mendelian diseases, all of which are located in the CDS of tumor suppressor genes (e.g., *TP53* germline mutations in LFS). Their MAFs are typically ≤ 0.01% and odds ratios (ORs) for cancer risk are as high as 20. Virtually all familial cancer syndromes are caused by these germline coding mutations. At the other end are common low-penetrance susceptibility alleles with MAFs >5% and ORs of 1.1-2.0.

The vast majority of hundreds if not thousands of these low-penetrance alleles are located in noncoding regions of the human genome, where our understanding of biological consequences and cancer causality is rudimentary^36, 37^. In comparison to other GWAS polymorphisms, the *TP53* noncoding variant (*rs78378222*[C]) is unique in cancer susceptibility, tissue-specificity, and pathogenicity. First, it is associated with an increased risk of multiple types of tumors with a MAF of 0.1-1.92% and ORs ranging from 1.39 to 4.55 (**Table S1**). Given the world population is estimated to reach 7.7 billion this year and assuming the MAF is 1.0 %, characterization of this variant will benefit ~150 million people worldwide at risk of cancer. Second, it, as a *TP53* variant, protects breast cancer, the most frequent LFS tumor type and increases the risk for soft-tissue sarcoma, the second most frequent LFS tumor type. It also increases the risk for prostate and colon tumors, two of the top four cancers with the highest incidences worldwide. Thus, this variant is both hypermorphic (an increase in normal gene function) in some tissues and hypomorphic (a partial loss of gene function) in others (**Fig. 3a**). Third, it is *bona fide* pathogenic in comparison to other reported germline variants potentially disrupting miRNA binding. There are multiple studies on polymorphisms disrupting miRNA coding sequences^38^ or polymorphisms on the 3′UTR miRNA binding sites of cancer genes like *BRCA1*^39^, *ESR1*^40^, and *KRAS*^41^. Among them, the SNP rs61764370 (also known as LCS6) in a let-7 miRNA complementary site within the 3′UTR of the *KRAS* oncogene^41^ is among the most studied. However, the clinical utility of the KRAS LCS6 variant has been significantly questioned since the initial publications showing it is associated with lung cancer^41, 42^, breast cancer^43^, and ovarian cancer ^42^. In the largest ever study with a total of 140,012 human subjects, the KRAS LCS6 variant did not increase the risk of ovarian cancer or breast cancer, regardless of the BRAC1/2 status; null results were also obtained for associations with overall survival for ovarian cancer, breast cancer, and all other previously reported associations for these cancers ^44^. Other than this *TP53* variant, few polymorphisms in cancer genes that potentially interact with miRNA have been validated using animal models. Therefore, ascertaining the pathogenic role of this *TP53* variant in this study facilitates our understanding of common low-penetrance noncoding cancer susceptibility loci.

On the basis of our findings and previous epidemiological studies, we compare cancer patients carrying rs78378222[C] to LFS patients. The rs78378222[C] differs from LFS mutants in several aspects: mutation sites (over 100 sites *versus* a single nucleotide), tumor spectrum (including breast cancer *versus* protective against breast cancer), penetrance (high *versus* moderate), and the size of the affected population (reported thousands *versus* potentially as many as 150 millions) (**Table 2**). Another difference is the impact of sex on cancer penetrance: the lifetime cancer risk of LFS patients is estimated to be 73% in males and nearly 100% in females, with the high risk of breast cancer accounting for the difference ^45^. The effect of sex on cancer penetrance may not exist or could even be reversed in patients carrying this variant, which confers risk for prostate cancer and protection for breast cancer. Finally, different miRNAs are involved in regulation of cancer susceptibility by LFS mutants and the noncoding variant. Relative hypomethylation of the promoter of *miR-34A*, a p53 target gene, is found in peripheral blood cells from LFS patients compared to that from patients with a wild-type *TP53*; hypermethylation of the *miR-34A* promoter is detected in tumors but not in histologically normal adjacent tissues from LFS patients^46^. These results support of a role of miR-34a in *inter-individual* cancer susceptibility, yet the causative relationship has not been investigated^46^. The present work, on the other hand, offers an example of miRNAs that are differentially expressed and modulate *intra-individual* tissue-specific cancer susceptibility. This is the first report showing that differential expression of miRNAs reduces the expression of a pathogenic genetic variant in one tissue yet increases its expression in another and contributes, at least partially, to tissue-specific cancer susceptibility within the same individual. Collectively, these differences, particularly that these patients with the noncoding variant are protected from breast cancer, argue against calling rs78378222[C] an LFS variant. In the clinical context of cancer clusters found in a family that fit the description (i.e., multiple family members developed various cancers associated with this variant as in **Table S1**), rs78378222 genotyping and a potential targeted cancer surveillance plan, different from the one for LFS ^47^, should be considered for those at risk.

**Table 2.** *TP53* germline coding mutations and the noncoding variant in cancer patients.

| Characteristic | LFS <sup>1</sup> | PAS <sup>2</sup> |
| --- | --- | --- |
| Altered loci | Coding (>100 mutation sites) | Noncoding (1 site) |
| Tumor spectrum | Breast cancer<br>Brain tumor (glioma)<br>Soft tissue sarcoma<br>Osteosarcoma<br>Adrenocortical carcinoma<br>Choroid plexus tumor<br>Rhabdomyosarcoma of embryonal anaplastic subtype | Protective against breast cancer<br>Brain tumor (glioma)<br>Soft tissue sarcoma<br>Prostate cancer<br>Colorectal adenoma<br>Uterine leiomyoma |
| Mean age at tumor onset | ~25 years | >>25 years |
| Population frequency | ~0.01% | ~2.0% |
| Sex effect on penetrance | Female > male | ? |
| Epigenetic regulation in susceptibility | miR-34a | miR-382 & miR-325 |

<sup>1</sup> The cancer spectrum in LFS was determined by comparing the incidence in LFS patients with that in the general population with similar demographics. Cancers like colorectal, prostate, and lung cancer and leukemia are found in LFS patients, yet they are not considered LFS-specific “core” cancer types because their incidence in LFS patients is not significantly higher than that in the general population. Li-Fraumeni-like (LFL) syndrome is similar to LFS, but does not meet the stringent classification criteria, i.e., the classic LFS definition established by Drs. Li and Fraumeni in 1988<sup>4</sup>. Birch<sup>30, 60</sup> and Eeles<sup>61</sup> have suggested two definitions of LFL, respectively. Regardless of the disease definition, *TP53* coding mutations are found in 20% to 40% of LFL families, and like LFS, breast cancer is the most frequent tumor type in LFL patients<sup>62</sup>. That patients with the *TP53* coding mutation have a later cancer onset than those with LFS mutations implies the influence of confounding genetic modifiers and the environment.
<sup>2</sup> The cancer spectrum was determined by genome-wide association studies (GWAS) of cancer cases and unaffected controls (**Table S1**). We did not investigate skin basal cell carcinoma, neuroblastoma, and esophageal squamous cell carcinoma in this study. The late tumor onset is suggested by our association study and mouse model data. Specific to breast cancer, the variant increases the risk in a Chinese population, but has no effect in a European population.

A strength of our study is the use of direct genotyping in determining cancer susceptibility rather than imputation. The *p53*^C^ allele delayed mammary tumorigenesis in two mouse models, and it is protective for breast cancer in this study with a Chinese population. In a previous study with a European population, there was no specific association between this variant and breast cancer^8^. This could be due to the inaccuracy of SNP imputation^8^, though we cannot rule out the possibility of ethnicity-specific differences in breast cancer protection. Another strength is that the biological plausibility of this variant in cancer risk and protection is extensively interrogated and ascertained using numerous mouse models and molecular cellular assays. However, our study has several limitations. First, the number of cases for soft tissue sarcoma is relatively small. Second, species-specific differences are noted: *p53*^+/C^ mice developed no glioma with *Pdgfb* overexpression in the SVZ. Third, there are many lingering questions to be addressed. What types of tumors do patients with *TP53*^C/C^ and children and adolescents with the variant develop? Are miRNAs behind the protective role of the *p53* variant against squamous cell carcinoma of head and neck^48^? Is p53 upregulated in some tissues to an extent so that carriers show moderate signs of ribosomopathies or the CHARGE syndrome, in which p53 is inappropriately activated^49, 50^? Further studies from population genetics and mouse models will reveal and corroborate the broader disease spectrum associated with this noncoding variant that may affect up to 150 million people worldwide.

## Experimental Methods

### Data reporting

No statistical methods were used to predetermine sample size.

### Patients

All recruited breast cancer and sarcoma patients, who are ethnic Han Chinese, were diagnosed at the Central Hospital of Wuhan, China. Patients who were suspected to have LFS or hereditary breast were excluded. Controls were unaffected (not diagnosed with any cancer) and were recruited at the same hospital with age (in 10-year intervals), sex, and ethnicity matched to the cancer cases. This study was approved by the Ethical and Scientific Committee of the Central Hospital of Wuhan, and all human subject research was performed in accordance with institutional, national, and Declaration of Helsinki requirements. The association of rs78378222[C] with the risk of cancer was determined using the Cochran-Armitage trend test.

### Genotyping

Peripheral blood samples were collected with informed consent, which was obtained from all study participants. Genotyping for rs78378222 was performed using the Taqman SNP Real-time PCR Assay (ThermoFisher Scientific, Waltham, MA). Less than 1% of the DNA samples failed the Taqman PCR genotyping and were genotyped using Sanger DNA sequencing. We arbitrarily selected 100 samples from the Taqman genotyping and directly sequenced the allele ─ DNA sequencing results were in 100% concordance with the Taqman results.

### Reagents

Antibodies against B220 (ab64100), Igκ light chain (ab190484), GFAP (ab7260), synaptophysin (ab32127), HA-tag (ab24779), PCNA (ab29), GD3 (ab11779), nestin (ab6142), olig2 (ab109186), E-cadherin (for IHC, ab76055), PCK (ab7753), p53 (for IHC, ab31333), phosphorylated-p53 Ser15 (for IHC, ab1431), vimentin (ab92547), and α-SMA (ab5694) were purchased from Abcam (Cambridge, MA). Antibodies against p53 (for western blotting [WB], CST2524), phosphorylated-p53 Ser15 (for WB, CST9248), BrdU (CST5292), GAPDH (CST5174), β-actin (CST4970) and α-tubulin (CST2125) were from Cell Signaling Technology (Danvers, MA). Antibodies against β-casein (SC-166684) and β-catenin (SC-7963) were from Santa Cruz Biotechnology (Dallas, TX). Azoxymethane (AOM), doxorubicin hydrochloride, and 3-methylcholanthrene were purchased from Sigma-Aldrich (St. Louis, MO). Dextran sulfate sodium (DSS; 36,000-50,000 MW) was purchased from MP Biomedicals (Santa Ana, CA). miRNA mimics and anti-miRNAs for miR-382, miR-325 and their controls (mirVana miRNA Mimic Negative Control #1 and Anti-miR Negative Control #1) were purchased from ThermoFisher Scientific (Waltham, MA). Lentiviral vectors expressing miRNA mimics and anti-miRNAs for miR-382, miR-325 and control miRNA were purchased from Sigma-Aldrich.

### Mice

All mice were housed in microisolator cages (maximum 5 per cage of same-sex animals) and maintained in climate/temperature- and photoperiod-controlled barrier rooms (22 ± 0.5°C, 12-12 h dark-light cycle) with unrestricted access to water and standard rodent diet (Teklad 2918, Harlan, Indianapolis, IN) unless otherwise indicated. The number of animals used in each experiment was estimated from published studies with statistically significant results. All mouse studies were performed in compliance with procedures approved by the Institutional Animal Care and Use Committees at Cleveland Clinic and Wuhan Central Hospital. The genetically modified PAS mutation knock-in mouse was generated by using zinc-finger nuclease technology (Fox Chase Cancer Center, Philadelphia, PA). The *p53*^1175A^ located in the *p53* PAS (AATAAA) was mutated to *p53*^1175C^ (AATACA), corresponding to human rs78378222. The targeting vector was electroporated into C57BL/6 embryonic stem cells and male chimaeras were bred with C57BL/6 WT females, and resulting F1 heterozygotes interbred to generate homozygotes in C57BL/6 background. The founders’ *p53* exons were sequenced in their entirety to confirm no mutations other than the targeted nucleotide. Genotyping was done using forward primer 5′-CTC CAG GGC CTA CTT TCC TT-3′ and reverse primer 5′-GGT AAG GAC CAT GTG CCA GT- 3′, followed by ApoI and EcoRV endonuclease double digestion after PCR and was confirmed by Sanger DNA sequencing. WT (*p53*^+/+^), heterozygous PAS (*p53*^+/C^), and homozygous PAS (*p53*^C/C^) mice were generated exclusively by breeding heterozygotes (*p53*^+/C^) in the C57BL/6 background (mice were born at the expected Mendelian ratio). When heterozygous PAS (*p53*^+/C^) mice were bred to *PyVT* (FVB/N-Tg(MMTV-PyVT)634Mul/J, Jackson Laboratory), *Erbb2* (FVB/N-Tg(MMTVneu)202Mul/J), or Hi-Myc (FVB-Tg(ARR2/Pbsn-MYC)7Key/Nci) mice in the FVB genetic background, F1 or F2 littermates were used for comparison and analyses (**Table S5)**. When p53^−/−^ and *p53*^+/−^ mice (B6.129S2-*Trp53^tm1Tyj^*/J) were used, they were bred to either C57BL/6 (**Fig. 1f**, **Fig. S2**) or a hybrid C57BL/6: FVB genetic background (with *Erbb2* mice, **Fig. 2f**; **Table S5**). We used both male and female mice in all experiments, except in mammary tumorigenesis studies where only females were used and in prostate tumorigenesis studies where only males were used. When mammary tumors were assessed, total measured tumor burden was calculated by summing up the volumes of all tumors developed in each animal.

### Histopathology and immunohistochemistry

For routine histological analysis, all studied tissues from mice were fixed in 10% neutral buffered formalin (Sigma-Aldrich), embedded in paraffin and evaluated by conventional haematoxylin and eosin (H&E) staining of serial sections cut at 5-μm thickness for pathological analyses. For IHC staining, deparaffinized and rehydrated sections were boiled in Na-citrate buffer (10 mM, pH 6.0) for 20 min for antigen retrieval. Sections were incubated with primary antibodies overnight at 4°C. Tissue sections were developed using the rabbit-specific HRP/DAB (ABC) Detection IHC Kit or Mouse on Mouse Polymer IHC Kit (Abcam). After haematoxylin counterstaining, the slides were dehydrated and mounted. For IHC, positive staining was indicated as a dark brown signal in the cells. Images were then acquired using an Olympus IX51 microscope and analyzed using cellSens Dimension software (Olympus, Center Valley, PA).

### Colon cancer and colitis model

Colon tumors were induced with AOM and DSS. Briefly, mice were injected intraperitoneally with 10 mg/kg AOM (Sigma-Aldrich) on day 1. Starting on day 7, mice were given drinking water containing 2% DSS for 7 days. Animals were then given untreated, regular drinking water. At 8 weeks after AOM administration, mice were sacrificed, colons were collected, and tumors were counted and measured under a dissecting microscope. To induce colitis, age- and gender-matched WT and mutant mice were given drinking water containing 2% DSS for 7 days, followed by regular drinking water and housed for up to 21 days. Tissue and whole blood were collected at day 0 (before DSS treatment), day 7, day 14 and day 21 for analysis. Colon histopathology was examined on H&E-stained sections. A complete blood count was obtained using Cell-Dyn 3500 Hematology Analyzer (Abbott, Abbott Park, IL).

### Retrovirus production

293GP cells (Clontech, Mountain View, CA) were seeded to 70% confluence in DMEM medium containing 10% fetal bovine serum (FBS). PDGFB-IRES-EGFP-expressing pQCXIX vector was reported previously^33^. This viral plasmid and a plasmid expressing VSVG were mixed with lipofectamine 2000 (ThermoFisher) according to manufacturer’s instructions and were added to the cell culture. After 4-6 h incubation, the medium was changed using fresh DMEM medium containing 10% FBS and then collected 48 h later. After filtration, the medium was centrifuged at 30,000*g* for 2.5 h at 4°C. The pellets were re-suspended in serum-free DMEM, and aliquots were frozen and stored at −80°C until use. The viral titer was then determined by infecting cells with serial dilutions of medium containing retrovirus in 10-fold increments. EGFP-expressing cells were then counted 48 h later and viral titer was determined by the lowest dilution that gave rise to EGFP-expressing cells.

### Intracerebral stereotaxic injection

The glioma model was generated as described previously by injecting retrovirus expressing *Pdgfb* to the mouse SVZ^33^. The mice were anesthetized with an intraperitoneal injection of ketamine/xylazine (100 mg/kg and 10 mg/kg respectively) and placed in a stereotaxic frame. An approximately 1 cm incision was made in the midline of the scalp to expose the bregma, and stereotaxic coordinates were determined. A burr hole was drilled through the skull, and a Hamilton (Hamilton Company, Reno, NV) syringe containing the retrovirus was then inserted into the brain. For targeting the dorsal lateral corner of the SVZ, we used the coordinates of 1 mm + 1 mm + 2.1 mm^33^, with the bregma as the reference. A volume of 1 µl was injected at 0.2 µL per minute. At the end of the injection, the needle was slowly retracted. The skin was sealed with Gluture Topical Tissue Adhesive (ThermoFisher). Following surgery, the mice were placed on a warming pad to recover. In the immediate post-op period, the animals were continuously monitored until fully awake. During the recovery period, animals were re-examined at 12 and 24 h post-op. After the first 24 h post-op, the observation schedule became once daily with weight, appearance, and behavior being monitored. For transplantation experiments, primary tumor cells from *p53*^C/C^ mice were prepared as reported ^51^ and injected into the SVZ of WT syngeneic recipient C57Bl/6 mice at a flow rate of 0.2 µl per minute. To downregulate the expression of miR-382 in the brain of *p53*^C/C^ mice, retrovirus carrying both *Pdgfb*^33^ and the Sigma-Aldrich MISSION miRNA Inhibitor (anti-miR-382) were injected into the mouse SVZ. The control inhibitor was Sigma-Aldrich MISSION miRNA Inhibitor Negative Control #1 (anti-ath-miR416; ath-miR416 is a miRNA from *Arabidopsis thaliana* with no homology to human and mouse miRNA or other gene sequences).

### Intraductal injection of mammary gland

Intraductal injections were performed as described^52–54^. Briefly, mice were anesthetized using ketamine/xylazine (100 mg/kg and 10 mg/kg respectively), and hair was removed in the nipple area with a commercial hair removal cream. Eighteen microliters of high-titer lentivirus expressing miR-382 mixed with 2 μL 0.2% Evans blue dye (Sigma-Aldrich) in PBS was injected in the third mammary gland using a 34-gauge needle (Hamilton). The titer of the viral particles expressing miR-382 was 9.1 × 10^8^ transducing units (TU)/mL with TUs measured by p24 antigen ELISA assays. For the control, the same amount (TUs) of control viral particles was used. Mice were handled in a biological safety cabinet under a stereoscope.

### Sarcoma development model

To induce sarcoma tumorigenesis in mice, 8- to 10-week-old body weight- and gender-matched mice were treated by intramuscular injection of 3- methylcholanthrene (MCA, Sigma-Aldrich) mixed with corn oil (5mg/ml) in the right hind leg. Mice were observed weekly for the development of intramuscular tumors for up to 200 days, and tumors were harvested for histological analysis.

### Survival and tumor burden studies

In accordance with IACUC guidelines, animals were allowed to live until humane endpoints were reached. Signs of terminal brain, mammary, and other tumor burden included the following: weight loss of 20%; peri-orbital hemorrhages; papilledema; epistaxis (nose bleeds); seizures; decreased alertness; impaired motor function; and/or impaired ability to feed secondary to decreased motor function, paresis or coma; changes in posture or ambulation ─ tense, stiff gait, ataxia, avoidance/inability to bear weight for 48 h, difficulty walking, inability to maintain upright position; restlessness; pacing; lethargy; tumor necrosis; systemic infection; signs of moderate to severe pain or distress; severe anemia; respiratory distress; or severe bleeding.

### Whole mount of mouse mammary gland

For whole-mount analysis, the fourth inguinal mammary glands were used by dissecting the inguinal mammary gland and fixing it on a microscope slide. The gland was immediately placed on a slide with Carnoy’s solution; blunt tweezers were used to gently configure the gland to its native orientation. The gland was fixed overnight before being subjected to re-hydration and carmine red staining as described ^55, 56^. The stained glands were then flattened, dehydrated, cleared in xylene, and mounted for microscopy.

### Western blotting analysis

Cell and tissue lysates were denatured in Laemmli sample buffer (Bio-Rad, Hercules, CA) and resolved by Tris-glycine SDS-PAGE (4-20% polyacrylamide, Mini-PROTEAN Precast Gels, Bio-Rad). After transfer to polyvinyl difluoride membrane, the membranes were probed with primary antibodies, followed by incubation with horseradish peroxidase-conjugated secondary antibodies for detection with Pierce ECL Western Blotting Substrate (ThermoFisher). Immunoblots were performed at least three times and representative blots reported. All intensity quantification for Western blot was performed using ImageJ software (Dr. Wayne Rasband, NIH, Bethesda, MD). The relative ratio of p53 with GAPDH, β-actin, or NPT-II as a reference was determined by calculating the relative density of the p53 bands normalized to that of the references. The relative ratio indicates the mean values of 3 independent blots in human cell experiments.

### RNA preparation and qPCR

To measure gene expression, total RNA was extracted from cultured cell lines or tissues and tumors dissected from mice using Trizol (Invitrogen) and run on a Synergy HTX Multi-Mode Microplate Reader (BioTek, Winooski, VT) to ensure that the A260/A280 ratio was in the range of 1.8-2.2 and the rRNA ratio (28S/18S) was >0.9. RNA was reverse transcribed with the iScript Reverse Transcription Supermix kit (Bio-Rad). Real-time PCR was performed using the iQ SYBR Green Supermix (Bio-Rad) and CFX96 Real-Time PCR detection system (Bio-Rad). The primers used for real-time PCR were designed based on the Universal Probe Library (Roche, Roche Life Science, Pleasanton, CA, USA) with GAPDH as reference. Real-time detection of miRNA expression was performed by using MystiCq miRNA qPCR Assay Primer (Sigma-Aldrich), using Universal Mouse Reference RNA and U6 as reference for mouse and human cells, respectively.

### FACS analysis of apoptosis and EGFP/RFP expression

Cells were plated in 6-well plates at a density of 2 × 10^5^ cells per well and cultured in Dulbecco’s modified Eagle’s medium (DMEM) and 10% FBS with penicillin and streptomycin until 80% confluence before treatment. Once confluence was reached, cells were collected and incubated with Alexa Fluor 488 Annexin V and propidium iodide using the Alexa Fluor 488 Annexin V/Dead Cell Apoptosis Kit (ThermoFisher). Cells were analyzed on a flow cytometer (MACSQuant Analyzer 10, Miltenyi Biotec, Bergisch Gladbach, Germany), and data were processed with the MACSQuantify software (Miltenyi Biotec). To detect EGFP/RFP expression, cells were co-transfected with EGFP/RFP-expressing plasmids containing the WT or the alternative PAS *TP53* 3′UTR (**Fig. 3**), or co-transfected EGFP-expressing plasmids containing the WT or the alternative PAS *TP53* 3′UTR with a RFP-expressing plasmid with WT *SV40* 3′UTR as transfection control (**Fig. S9**), using Lipofectamine LTX (ThermoFisher) according to the manufacturers’ protocols.

### Cell culture

Human glioblastoma multiforme cell line U-87 (HTB-14, ATCC, Manassas, VA), colon cancer cell line HCT116 (ATCC CCL-247), HCT116 (p53^−/−^, courtesy of Dr. Bert Vogelstein), cervical cancer cell line HeLa (ATCC CCL-2), non-small cell lung cancer cell line H1299 (ATCC CRL-5803), breast cancer cell line MDA-MB-231 (ATCC CRM-HTB-26), MCF7 (ATCC HTB-22), embryonic kidney cell line HEK293(ATCC CRL-1573) and 293GP cells (Clontech) were cultured in high glucose (5g/l) DMEM and 10% FBS with penicillin and streptomycin at 37°C in 5% CO_2_. Human breast epithelial cell line MCF10a (ATCC CRL- 10317) was cultured in complete MEGM media (Lonza, Walkersville, MD) with 100 ng/ml cholera toxin at 37°C in 5% CO_2_. Cell lines from ATCC and Clontech were authenticated using short-tandem repeat profiling. All cells were tested *Mycoplasma* free and used for less than 10 passages upon acquisition. We performed SNP genotyping for all these human cell lines and found that none of them carried rs78378222[C].

### Senescence-associated β-galactosidase assay

MEFs were prepared from E13.5 embryos and senescence of MEFs was determined by using the Senescence β-Galactosidase Staining Kit (Cell Signaling) according to the manufacturer’s protocol^57^. MEFs were plated at 2 × 10^5^ cells per well in a 6-well plate and grown overnight. After 24 h, cells were rinsed with sterile PBS, fixed with 0.5% glutaraldehyde at room temperature for 15 min, and then rinsed with sterile PBS. Cells were stained with fresh senescence-associated β-galactosidase staining solution (Cell Signaling) at 37°C for 24 h. The staining solution was then removed, and cells were stored in PBS at room temperature. Senescent cells show marked perinuclear blue staining, whereas non-senescent cells do not exhibit this stain.

### miRNA targeting and luciferase assay

miRNA targeting prediction was performed using TargetScanS^58^ and PolymiRTS^59^. To determine p53 activity, cells were co-transferred with pNL(NLucP/p53-RE/Hygro) (Promega, Madison, WI) and the desired p53-expressing plasmids. A luciferase assay was then performed using the Nano-Glo Luciferase Assay System (Promega). Samples were then run on Synergy HTX Multi-Mode Microplate Reader (BioTek) to record the signal intensity. This pNL(NLucP/p53-RE/Hygro) plasmid contains several copies of a p53 response element (p53-RE) that drives transcription of a destabilized form of NanoLuc luciferase, an engineered small (23.3 kDa) luciferase fusion protein. The NlucP reporter consists of NanoLuc luciferase with a C-terminal fusion to PEST, a protein destabilization domain, which responds more quickly and with greater magnitude to changes in transcriptional activity than unmodified NanoLuc luciferase.

### RNA pull-down assays

5′end biotin-labeled *TP53* 3′UTR RNA fragments with different PAS and other mutations and miRNA of interest labeled with biotin at the 3′end were commercially synthesized (Sigma-Aldrich). H1299 cells were seeded one day before transfection in 10 cm tissue culture dish at 50% confluent. Twenty four hours later, cells were transfect with 5′end biotinylated *TP53* 3′UTR RNA fragments (final concentration of 100 nM) or 3′end biotinylated miRNA (final concentration of 100 nM), and/or a plasmid carrying a *TP53* 3′UTR (10 µg) according to manufacturer’s guideline. Twenty four hours post transfection, cells were lysed in 550 μl lysis buffer supplemented with protease inhibitors and RNase inhibitor, followed by incubation on ice for 10 min. The cell lysates were centrifuged at 4°C, 14,000 *g* for 10 min and the supernatant was subjected to Pierce™ Streptavidin Magnetic Beads in the presence of 2% RNase-free BSA (ThermoFisher Scientific) and 2% yeast tRNA (ThermoFisher Scientific) to block non-specific binding. After 2 hour of incubation, the beads were placed on a magnetic stand for 8 min and washed 4 times with 1 ml lysis buffer. Purified RNAs were extracted by RNAzol® RT (RN 190, Molecular Research Center; Cincinnati, OH) and subjected to reverse transcription and quantitative real-time PCR to determine the beads-bound miRNAs or 3’UTRs.

### Statistical and data analysis

The product-limit method of Kaplan and Meier was used for generating mouse survival or tumor-free survival curves, which were compared by using the log-rank (Mantel-Cox) test. An unpaired two-tailed Student’s t-test was performed for two-group comparisons. One-way analysis of variance (ANOVA) was performed for multiple group comparisons with one independent variable and two-way ANOVA for multiple group comparisons with two independent variables (genotype or treatment). Tumor multiplicity was compared using the Mann-Whitney *U* test. A *P* value less than 0.05 was considered statistically significant for all datasets. All statistical analyses were performed using SPSS 16 (IBM Analytics, Chicago, IL), GraphPad Prism 5 (GraphPad Software, La Jolla, CA), or Sigmastat 4 (Systat Software, London, UK).

## Acknowledgments

The authors are grateful to Dr. Cassandra Talerico, a salaried employee of the Cleveland Clinic, for editing the manuscript and providing critical comments and to Dr. James F Crish for retroviral generation and injection.

## Author contributions

Q.D., H.H., X.Y., Z.L., and Y.L. conceived the model, designed and carried out experiments, and wrote the manuscript. All authors contributed to research and data analyses and approved the final version of the manuscript. Correspondence should be addressed to Y.L. or Z.L.

## Data and materials availability

All data are available in the main text or the supplementary materials and methods. The *p53* mutant mouse line is available to any researcher upon request.

---

**Fig. S1.**
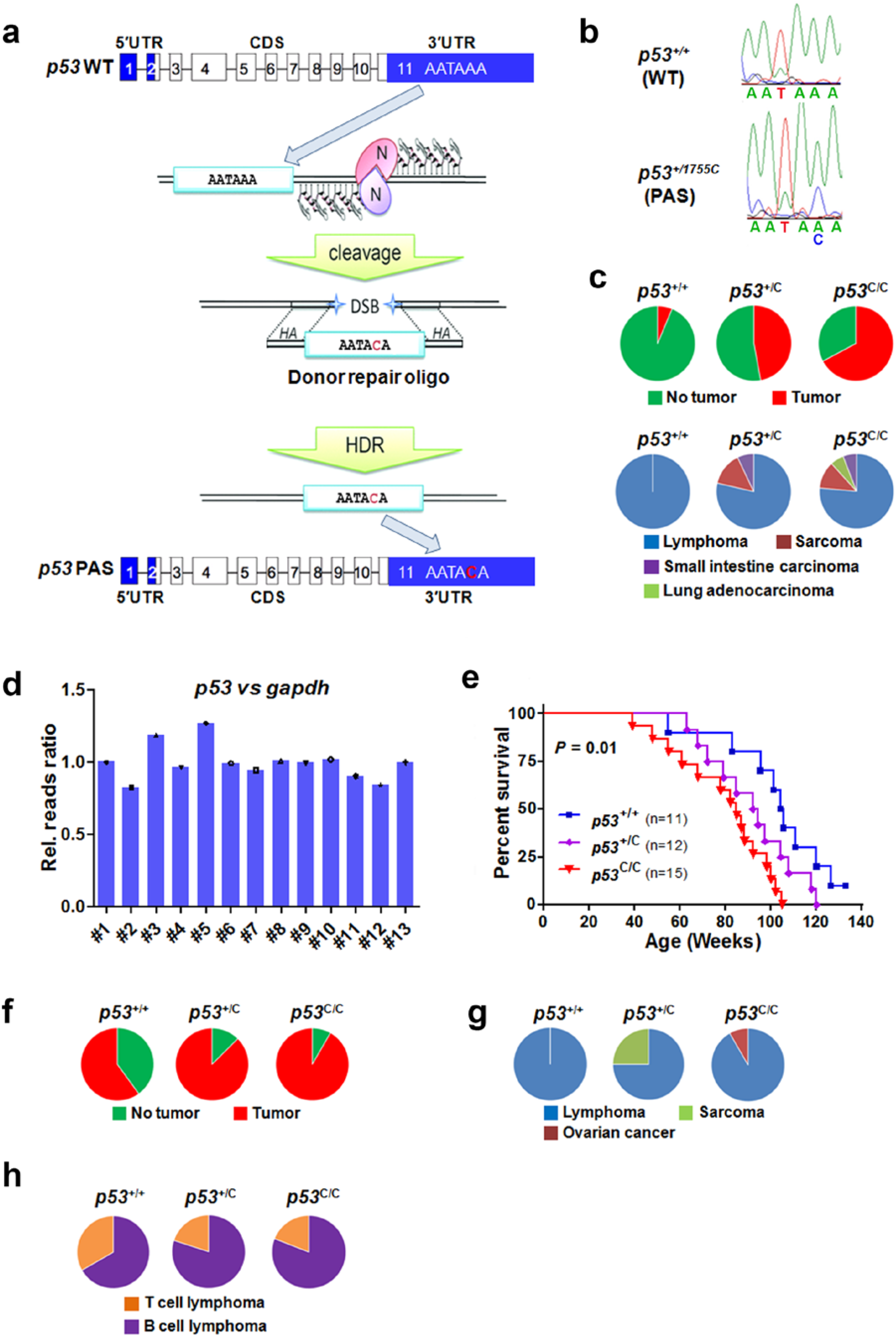
The *p53*^C^ allele accelerates spontaneous tumorigenesis and radiation-induced tumorigenesis in mice. (**a**) Schematic diagram showing the mutation at the endogenous *p53* gene polyadenylation signal (PAS) with zinger finger nuclease-mediated cleavage and homology-directed repair. Numbers in rectangles indicate exons of the *p53* gene; blue rectangles indicate untranslated regions (UTR) and clear rectangles indicate coding sequences. Lines connecting rectangles indicate introns in the *p53* gene. (**b**) Confirmation of the mutation introduction by DNA sequencing. (**c**) Proportions of *p53*^+/+^, *p53*^+/C^, and *p53*^C/C^ littermates that developed spontaneous tumors during their lifespan; tumor types were shown. (**d**)The relative read ratio (Rel. read ratio) for *p53* normalized to that of *Gapdh* in genomic DNA isolated from B cell lymphoma that developed in mutant mice, as determined by exome sequencing. (**e**) Survival of *p53*^+/+^, *p53*^+/C^, and *p53*^C/C^ littermates treated with a single dose of 4 Gy γ-irradiation at 6 weeks of age. *P* = 0.01 (log-rank test). (**f**) Proportions of mutant and WT littermates that developed tumors during their lifetimes after irradiation. (**g**) Cancer spectrum of irradiated mice. (**h**) Proportions of T cell lymphoma and B cell lymphoma in irradiated mice.

**Fig. S2.**
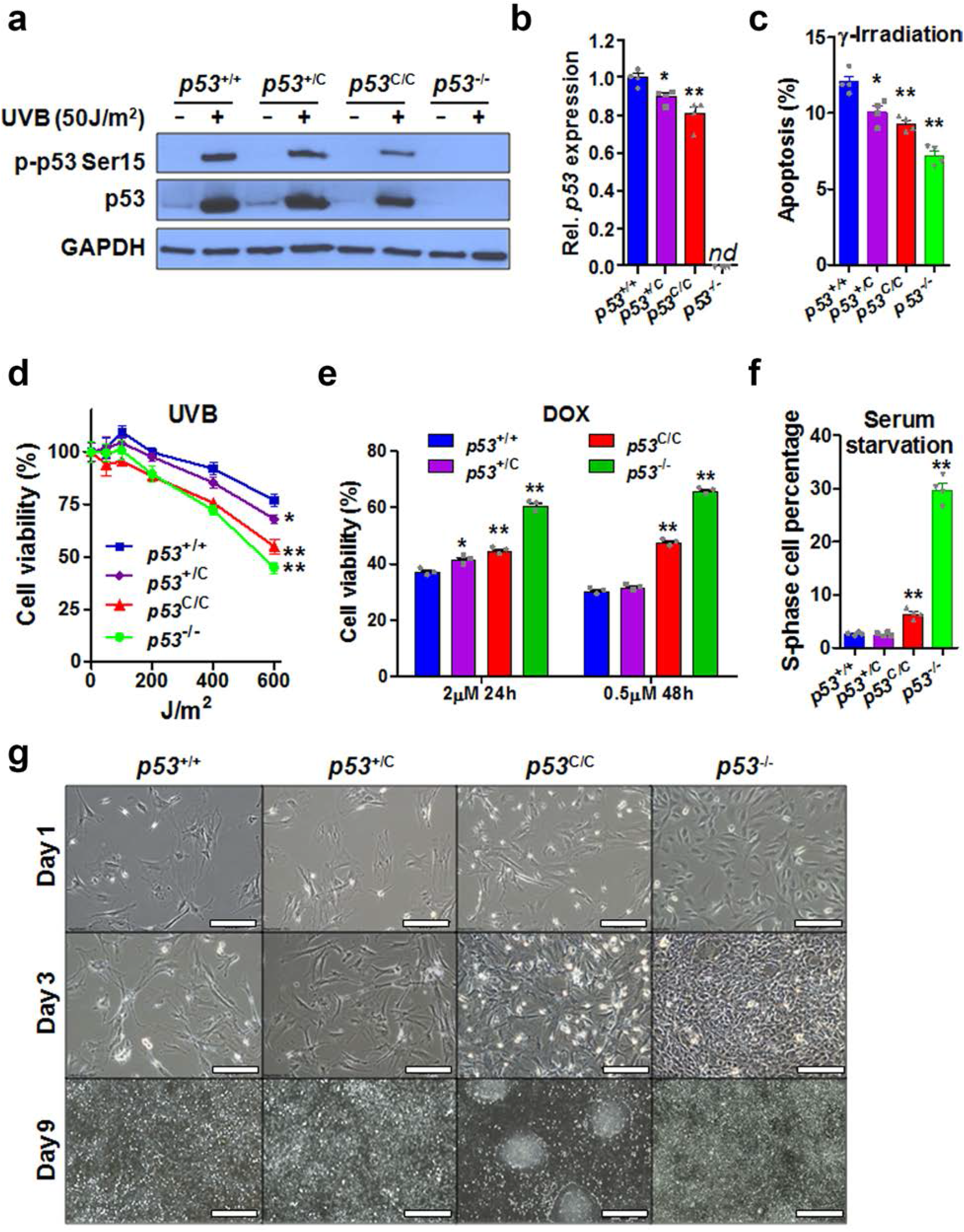
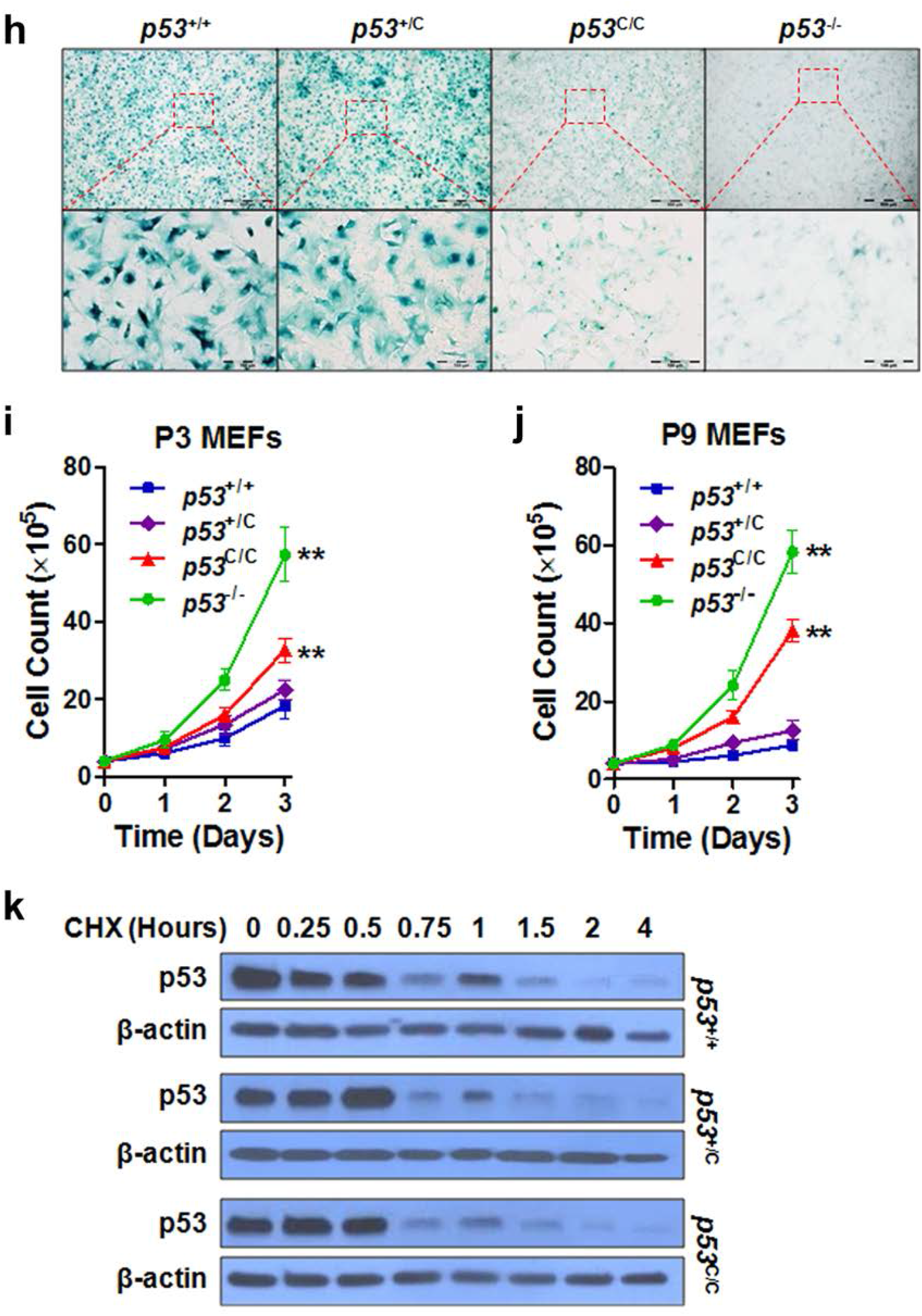
Reduced *p53* expression and function in mouse embryonic fibroblasts (MEFs) isolated from mice with the *p53*^C^ allele. *n* ≥ 3 mice for each genotype. Data were presented as mean ± s.e.m; student’s t-test was performed to compare with *p53*^+/+^ cells; * *P* < 0.05, **, *P* <0.01. (**a**) Western blotting analysis of p53 and phosphorylated p53 in primary MEFs isolated from mice with indicated genotypes. MEFs were treated with 50 J/m^2^ UVB and analyzed 6 h post-treatment. (**b**) Relative *p53* mRNA levels in WT and mutant MEFs as measured by qPCR. nd, not detectable. (**c**) Apoptosis analysis of MEFs treated with 10 Gy γ-irradiation. At 24 h post treatment, FACS was used to detect annexin V-FITC-positive cells. (**d**) Viability of primary MEFs treated with indicated dosage of UVB. Cells were analyzed 48 h post treatment using the MTT assay. (**e**) Viability of primary MEFs treated with 0.5 or 2 µM doxorubicin (DOX). Cells were analyzed 24 or 48 h post treatment using the MTT assay. (**f**) Fraction of serum-starved primary MEFs in S phase as determined by FACS detection of propidium iodide-stained cells. (**g**) Morphologically representative images of MEFs with indicated genotypes (passage 3) from day 1 to day 9. Scale bar, 200 μm; *n* = 3 independent MEF isolates (each with 3 technical repeats). (**h**) SA-β-gal staining of MEFs for the indicated genotypes (passage 9). Scale bars: 500 μm (top row), 100 μm (bottom row). (**i**, **j**) Quantitative analysis of MEF growth with indicated genotypes at passage 3 or 9. (**k**) Western blotting for p53 expression in MEFs treated with UVB and cycloheximide (CHX). Cells were treated with 200 J/m^2^ of UVB; 6 h later, cells were incubated with 100 µM CHX for up to 4 h before analyses.

**Fig. S3.**
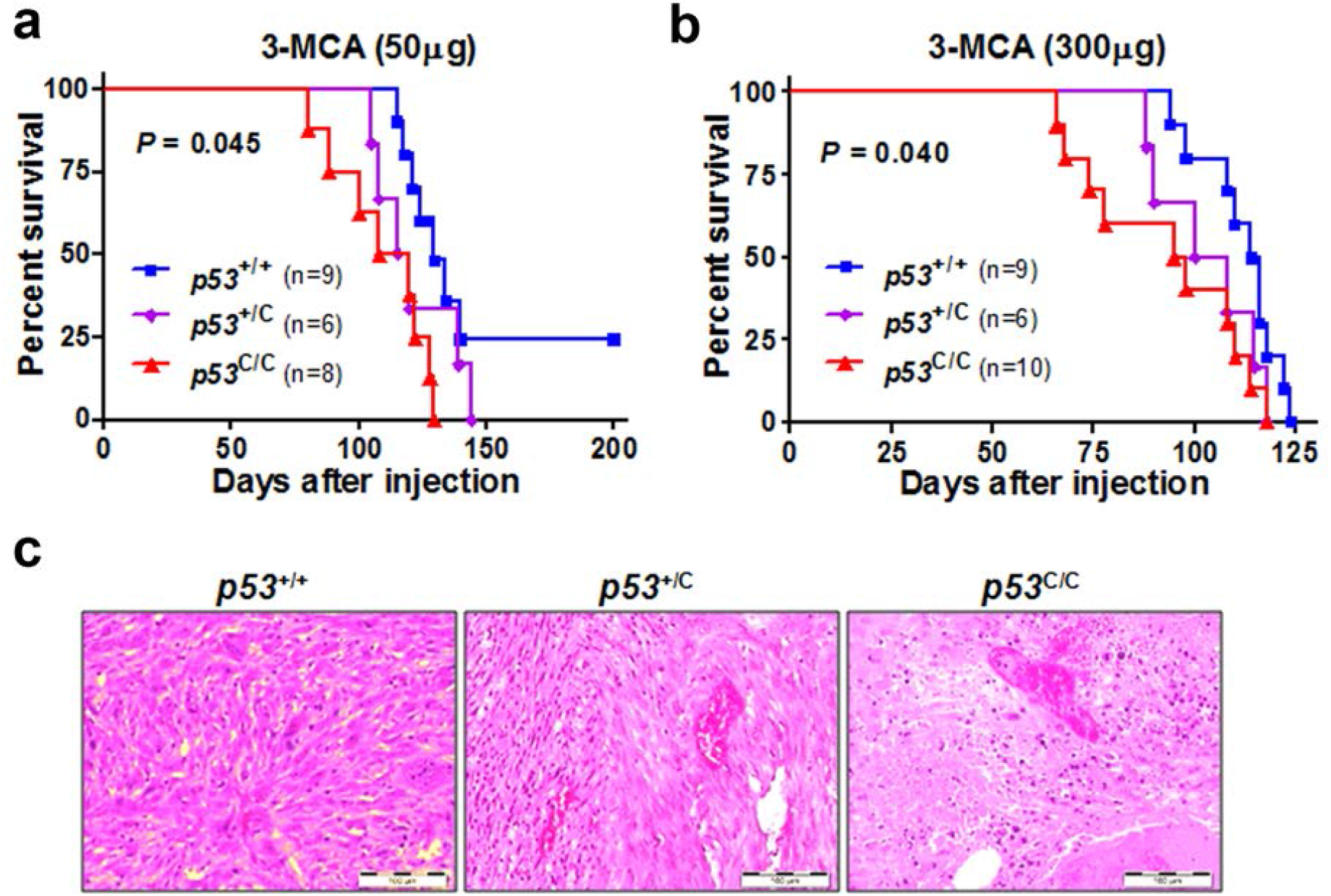
The *p53*^C^ allele enhances carcinogen-induced sarcoma development in mice. (**a, b**) Survival of mice injected with 50 μg (**a**) or 300 μg (**b**) of 3-methylcholanthrene (3-MCA) to skeletal muscle of left hind leg. *P* = 0.045 (**a**) and *P =* 0.04 (**b**). n = 6 to 10 mice per group. (**c**) Representative H&E staining for sarcoma in mice treated with 50 μg 3-MCA. Scale bar, 100 μm; *n* = 3 tumors (1 tumor per animal).

**Fig. S4.**
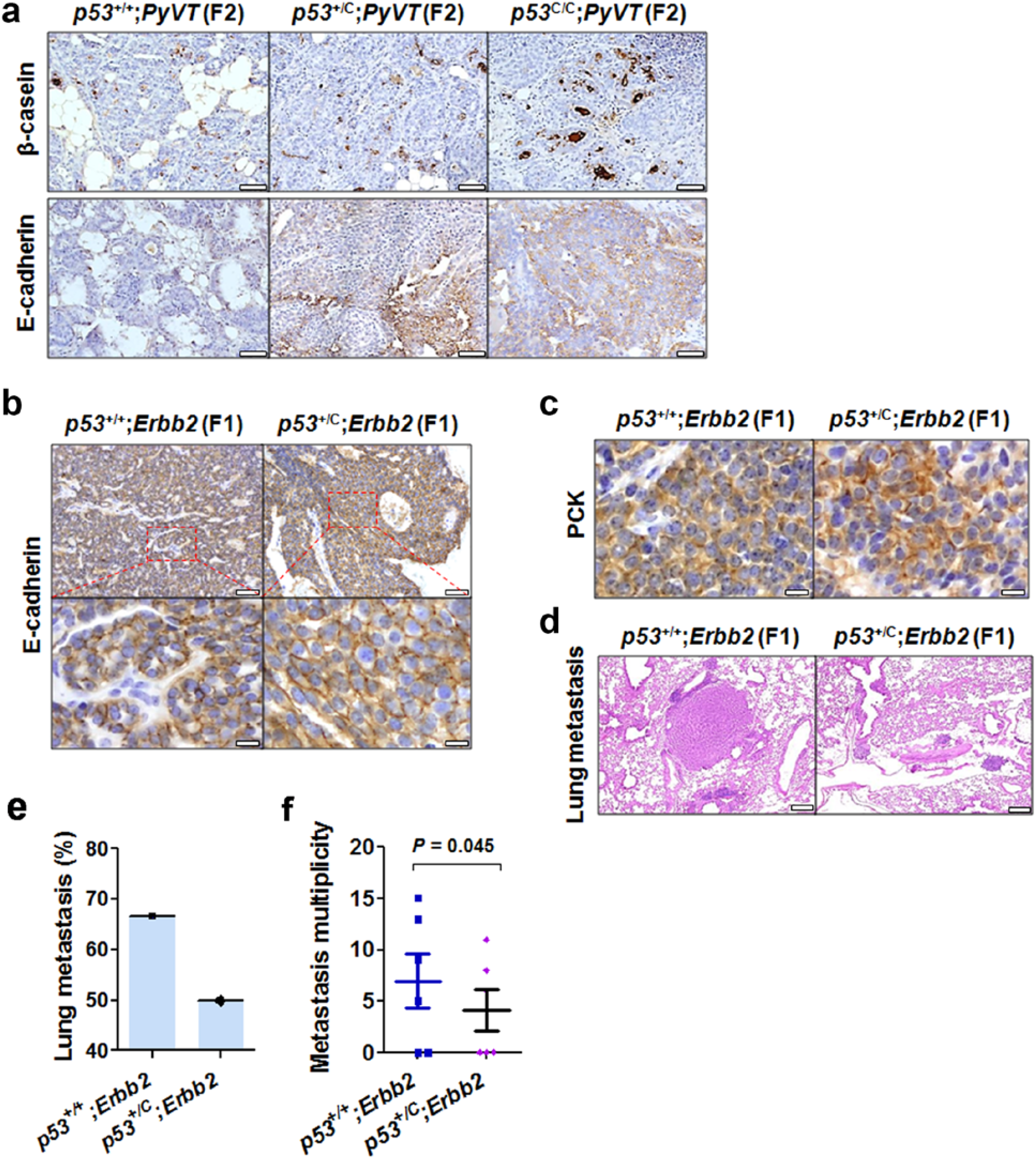
The *p53*^C^ allele delays mammary tumor progression and metastasis in mice. (**a**) Immunohistochemistry (IHC) staining of β-casein and E-cadherin in tumors from 12-week-old F2 *PyVT* females. *n* = 3 mice per group. Scale bar, 50 μm. (**b**) IHC staining for E-cadherin in mammary tumors from F1 *p53*^+/+^;*Erbb2* mice and *p53*^+/C^;*Erbb2* mice. Scale bar, 100 μm (top row), 20 μm (bottom row). *n* ≥ 3 tumors for each genotype (one tumor per animal). (**c**) IHC staining for the epithelial marker pan cytokeratin (PCK) in mammary tumors from F1 *p53*^+/+^;*Erbb2* and *p53*^+/C^;*Erbb2* mice. Scale bar, 20 μm. *n* ≥ 3 tumors for each genotype (one tumor per animal). (**d**) Representative H&E staining for lung metastasis arising from mammary tumors in *p53*^+/+^;*Erbb2* and *p53*^+/C^;*Erbb2* mice. Scale bar, 200 μm. (**e**) Percentage of animals with lung metastasis. (**f**) Lung metastasis multiplicity. *n* = 10 mice for each genotype.

**Fig. S5.**
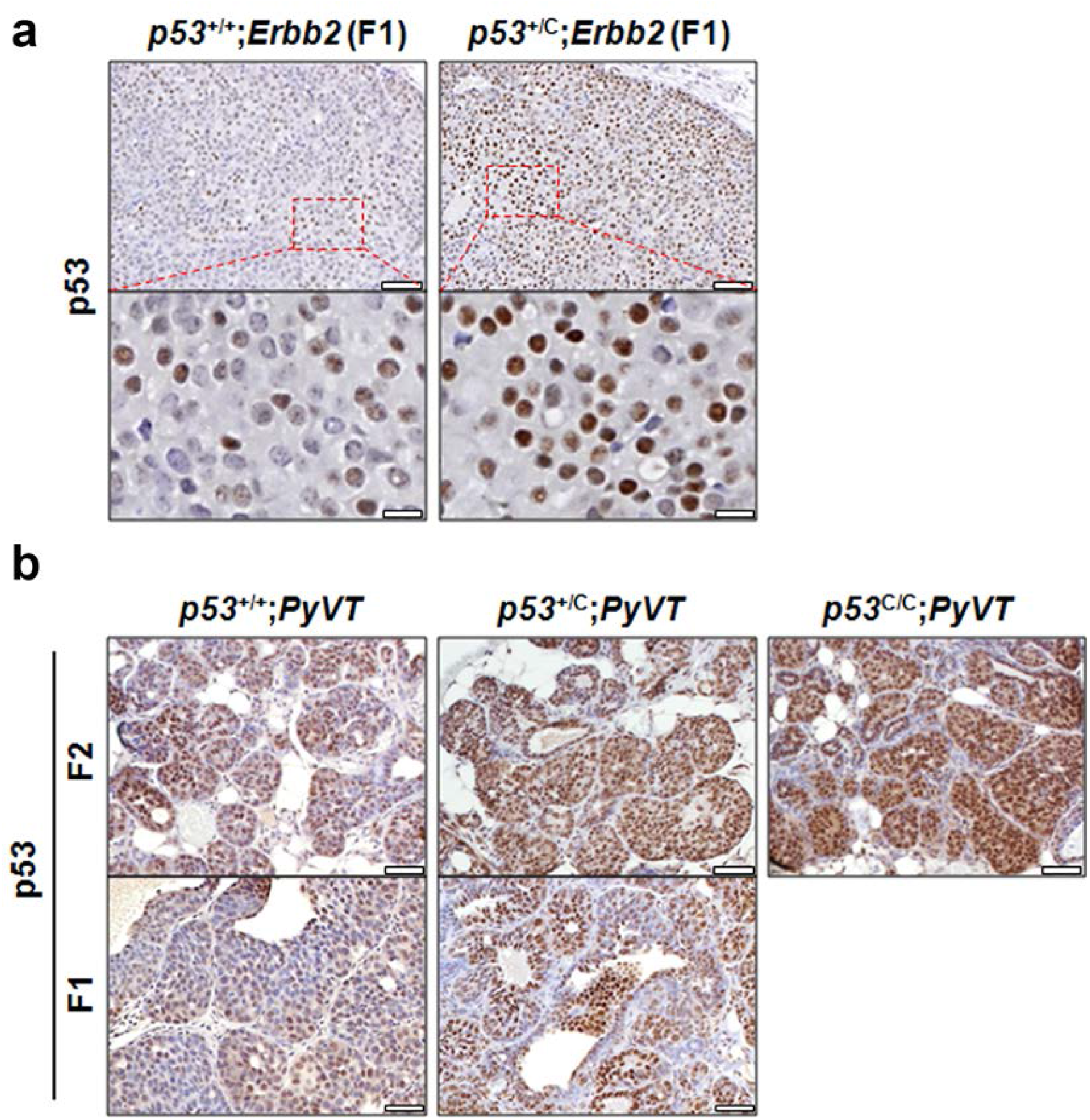
p53 protein levels in mammary tumors that developed in mice. (**a**) IHC staining for p53 protein levels in mammary tumors from *p53*^+/+^;*Erbb2* and *p53*^+/C^;*Erbb2* mice. Scale bar, 100 μm (top row), 20 μm (bottom row). *n* ≥ 3 tumors for each genotype. (**b**) IHC staining for p53 protein levels in mammary tumors from *p53*^+/+^;*PyVT*, *p53*^+/C^;*PyVT* and *p53*^C/C^;*PyVT* mice, either F2 or F1. Scale bar, 50 μm. *n* ≥ 3 tumors for each genotype.

**Fig. S6.**
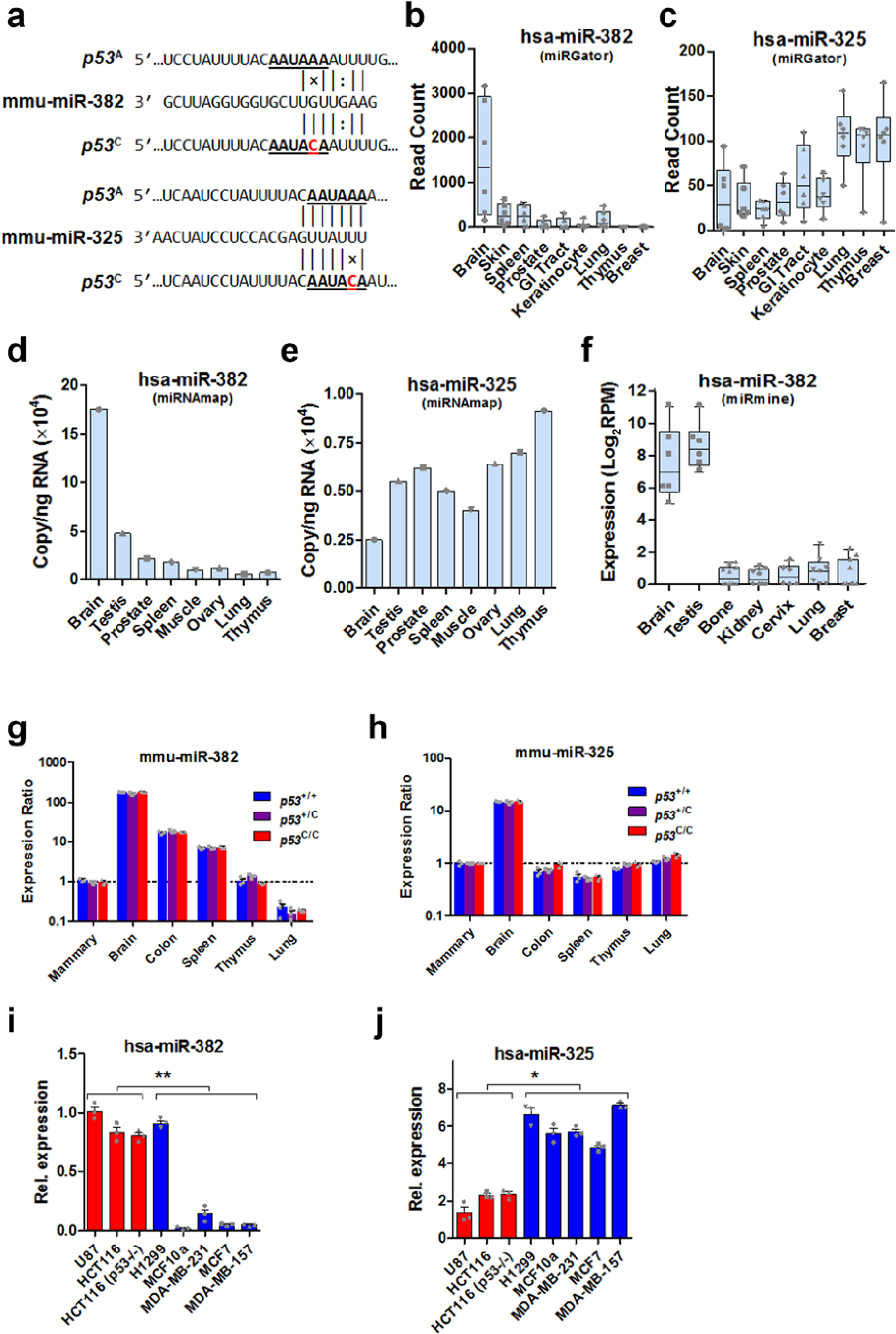
miR-325 and miR-382 expression is tissue-specific. (**a**) Schematic diagram showing the creation of a miR-382 targeting site and the disruption of a miR-325 targeting site in mouse *p53* 3′UTR due to the *p53*^C^ allele. (**b** and **c**) Expression levels (read count) of miR-382 (**b**) and miR-325 (**c**) in different human tissues by deep sequencing. Data were obtained from the online database miRGator. (**d** and **e**) Expression level (copy number) of miR-382 (**d**) and miR-325 (**e**) in indicated human tissues as determined by deep sequencing. Data were extracted from the online database miRNAmap, which contained no data on the expression of miR-382 and miR- 325 in breast tissue. (**f**) Expression level of miR-382 in indicated human tissues. Data were extracted from the miRmine database, which contained no miR-325 expression data; log_2_RPM, represents the logarithm of the number of miRNA reads per million total RNA reads to the base 2. (**g**) Relative expression of miR-382 in indicated tissues from *p53*^+/+^, *p53*^+/C^ and *p53*^C/C^ mice measured by qPCR. *n* = 3 mice, each performed with technical triplicates. (**h**) Relative expression of miR-325 in indicated tissues from *p53*^+/+^, *p53*^+/C^ and *p53*^C/C^ mice measured by qPCR. *n* = 3 mice. Please note the differences in scales between **g** and **h**. (**i** and **l**) qPCR analysis for miR-382 (**i**) and miR-325 (**j**) levels in human cancer cell lines. Rel. expression: Relative expression of miRNAs normalized to U6 RNA. Red columns indicate cell lines established from human glioma and colon cancers, and blue columns indicate cell lines established from human breast and lung cancers. *n* = 3 independent biological replications with technical triplicates. * *P* < 0.05, ** *P* < 0.01 (one-way ANOVA).

**Fig. S7.**
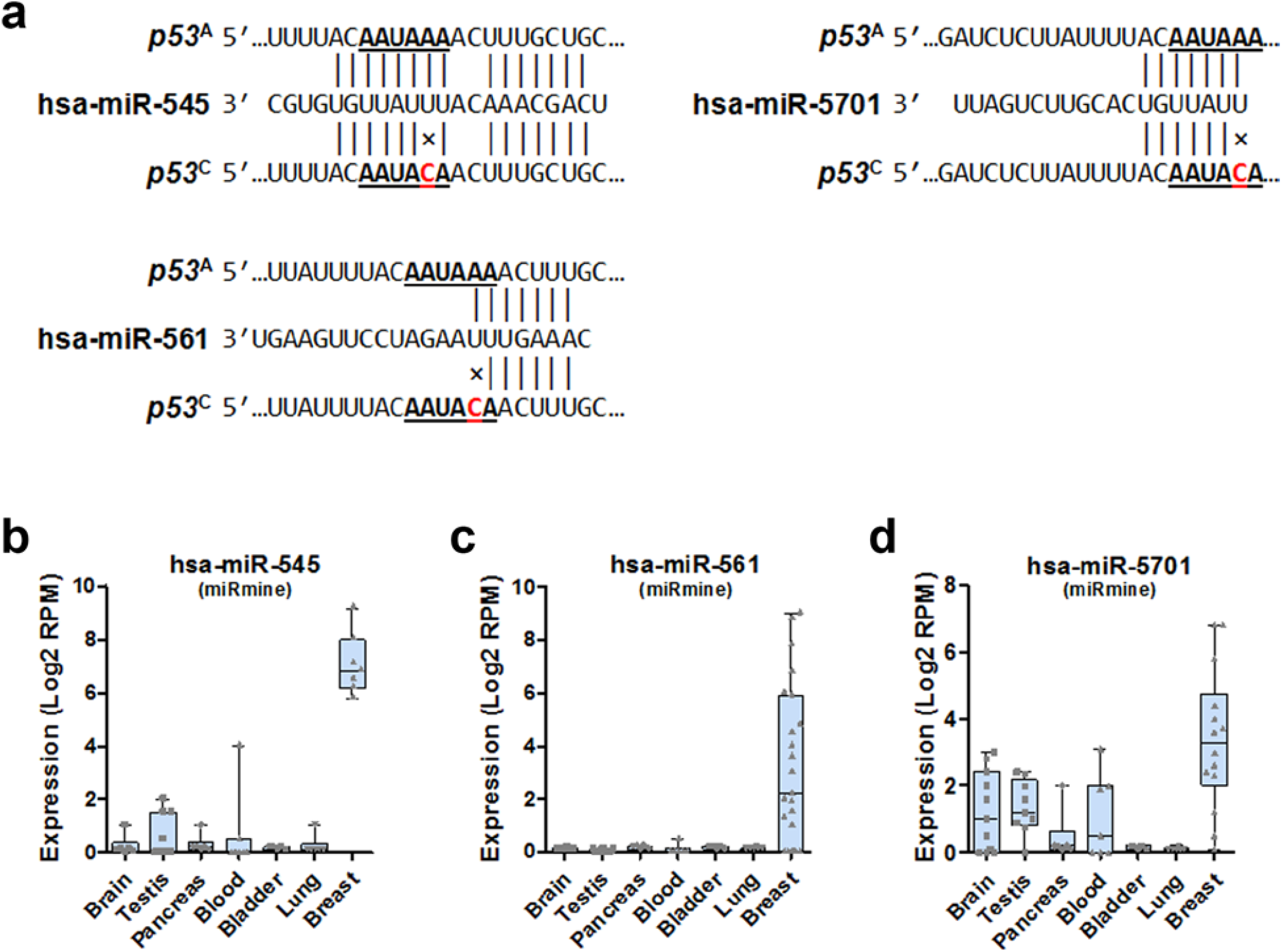
The expression of other miRNAs targeting the WT *TP53* 3′UTR. (**a**) Schematic diagram showing the disruption of miR-545, miR-561, and miR-5701 targeting sites in human *TP53* 3′UTR due to the PAS variant. (**b-d**) Expression level of miR-545 (**b**), miR-561 (**c**) and miR-5701 (**d**) in indicated human tissues as determined by deep sequencing. Data were extracted from the online database miRmine.

**Fig. S8.**
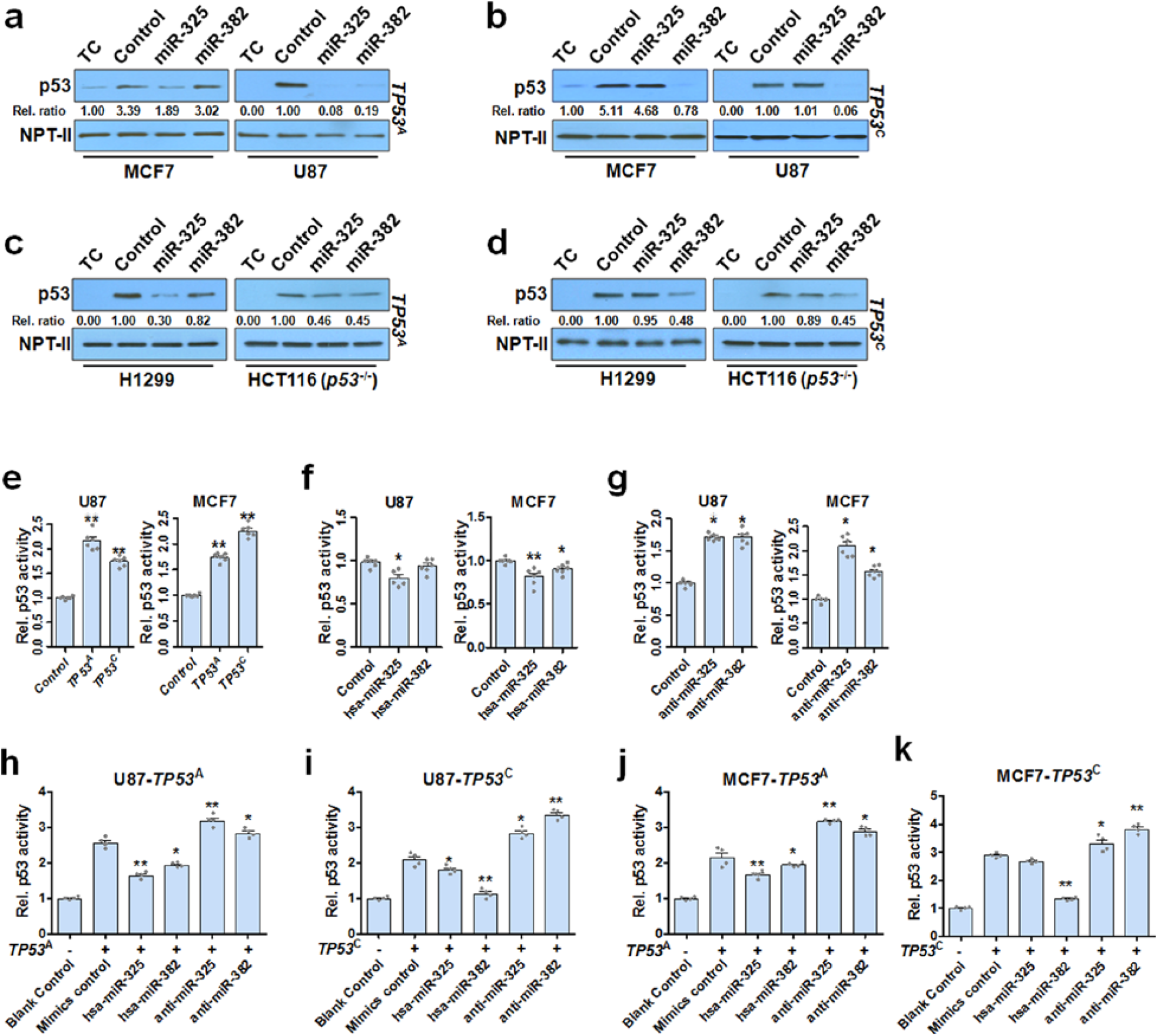
miRNA suppression of p53 expression and p53-driven luciferase activity. **a-d**, Western blotting analysis for p53 expression in several cell lines transfected with a p53*-* expressing plasmid with WT or the variant 3′UTR (*TP53*^A^ or *TP53*^C^) and miR-325 or miR-382 mimics. Control, transfection with the *TP53*^A^ or *TP53*^C^ with miRNA mimics negative control. Rel. ratio: the relative signal densities of p53 as referenced to that of NPT-II. *n* ≥ 3 experiments for each transfection and western blotting. (**a**) MCF7 and U87 cells transfected *TP53*^A^ and miR- 325 or miR-382. (**b**) MCF7 and U87 cells transfected with *TP53*^C^ and miR-325 or miR-382. (**c**) H1299 and HCT116 (*TP53*^-/-^) cells transfected with *TP53*^A^ and miR-325 or miR-382. (**d**) H1299 and HCT116 (*TP53*^-/-^) cells transfected with *TP53*^C^ and miR-325 or miR-382. TC = transfection control. **e**-**k**, NanoLuc luciferase assays for p53 transcriptional activity in U87 and MCF7 cells. All cells were transfected with pNL(NLucP/p53-RE/Hygro) (Promega), along with an exogenous p53-expressing plasmid with a native 3′UTR (*TP53*^A^) or a 3′UTR with an alternative PAS (*TP53*^C^) (**e**), miRNA mimics (**f**), or anti-miRNAs (**g**), or a combination of p53 expression plasmid and miRNA mimics (or anti-miRNAs) (**h**-**k**). Rel. p53 activity: relative p53 activity as determined by relative luminance units. Either endogenous or exogenous p53 drives NlucP expression. Data were presented as mean ±s.e.m; *n* = 3-6 for each group and *n* = 3 for biological replications. * *P* < 0.05, **, *P* < 0.01.

**Fig. S9.**
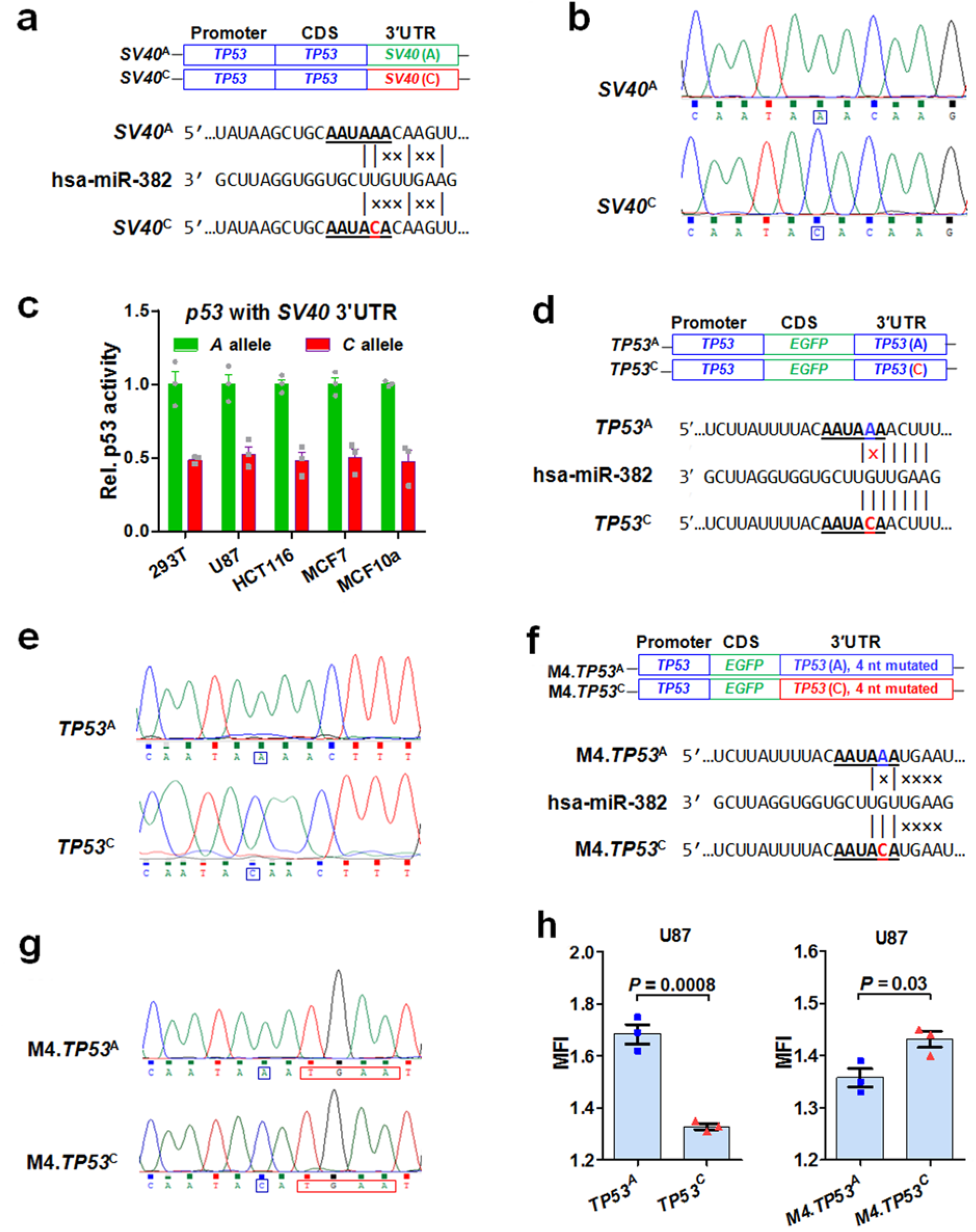
The integrity of the miR-382 binding sites in miR-382-mediated *TP53*^C^ suppression. (**a**) Schematic diagram and (b) Sanger DNA sequencing showing p53 expression plasmids with SV40 3′UTR WT PAS [*SV40* (A)] or the PAS variant [*SV40* (C)] and the absence of the miR-382 targeting site in these sequences. (**c**) NanoLuc luciferase assay for p53 transcriptional activity in indicated cell lines transfected with plasmids shown in (**a**). Rel. p53 activity: relative p53 activity as determined by relative luminance units, each normalized to cells with the *A* allele. Data were presented as mean + s.e.m; *n* = 3 for each group. (**c**) Schematic diagram showing EGFP expression plasmids with *p53* 3′UTR with WT PAS [*TP53* (A)] or the variant PAS [(*TP53* (C)] and the presence of miR-382 targeting sites (left). The mutation of 4 nucleotides (M4) in *TP53* 3′UTR with WT PAS [M4.*TP53* (A)] or a PAS variant [M4.*TP53* (C)] eliminated the miR-382 targeting site (right). (**d**) Schematic diagram and (**e**) sequencing showing EGFP expression plasmids with WT *p53* 3′UTR (*TP53*^A^) or the *TP53* 3′UTR variant (*TP53*^C^) and the presence of miR-382 targeting sites. (**f, g**) The mutation of 4 nucleotides (M4) in WT *TP53* 3′UTR (M4.*TP53*^A^) or the *TP53* 3′UTR variant (M4.*TP53*^C^) eliminated the miR-382 targeting site. (**h**) EGFP expression in U87 cells transfected with the 4 plasmids constructed in (**d** and **f**). An RFP expression plasmid with the native *SV40* 3′UTR with no miR-382 sites was co-transfected as the control. Y axis denotes the MFI (mean fluorescence intensity) for EGFP normalized to that of RFP. *n* = 3 biological triplicates and *n* = 3 technical triplicates.

**Fig. S10.**
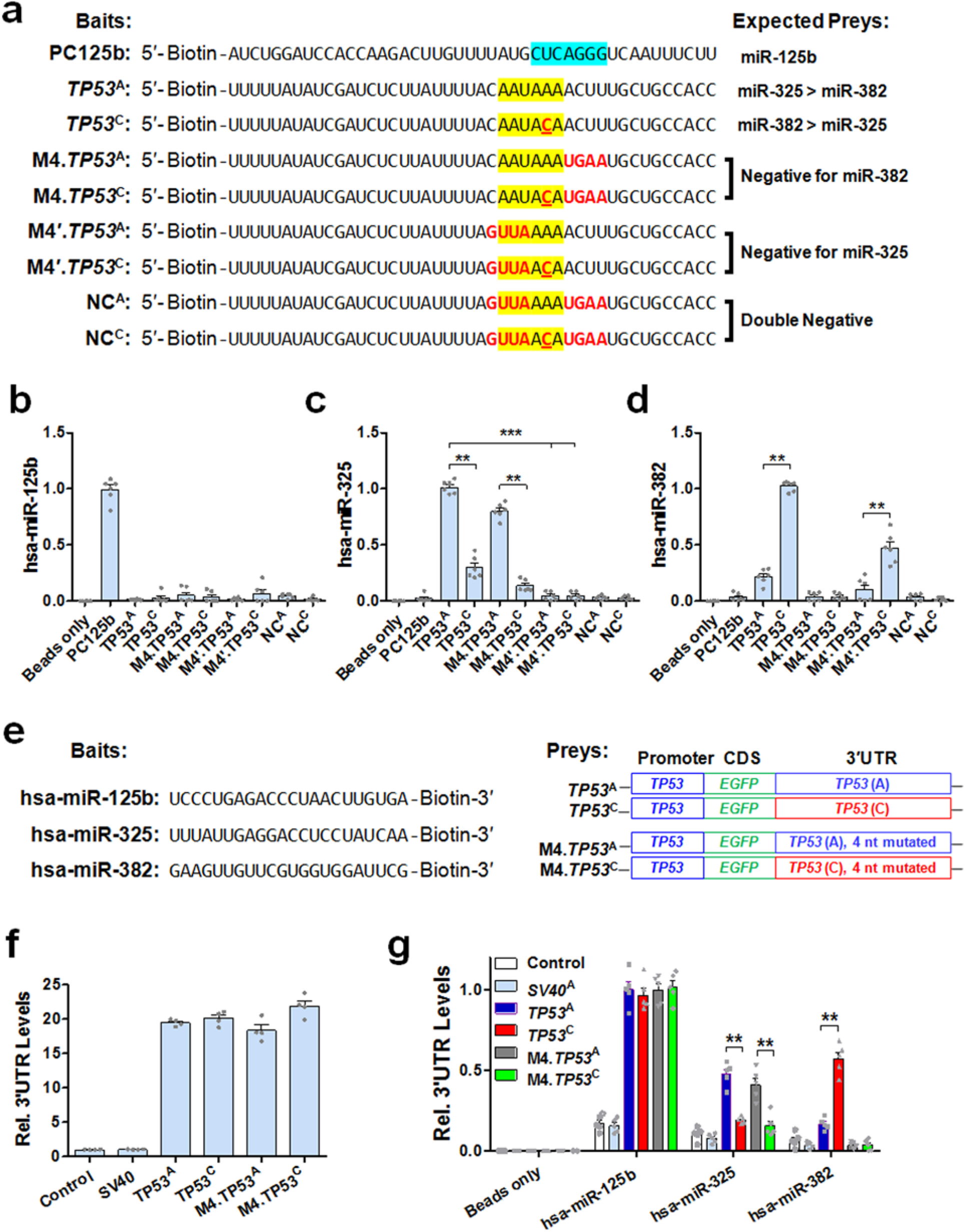
Direct binding of *TP53* 3′UTR and miRNAs using pull-down assays. (**a**) Schematic diagram showing the synthesized 5′end biotin-labeled *TP53* 3’UTR RNA fragments. Blue block, sequence complementary to miR-125b seed sequence; yellow block, PAS; bold letters in red, mutated nucleotides to remove miR-325 or miR-382 binding sites. PC125b: RNA fragment of *TP53* 3′UTR containing the miR-125b binding site as a positive control; *TP53*^A^: RNA fragment of *TP53* 3′UTR containing the WT PAS (*A* allele); *TP53*^C^: same as *TP53*^A^ except the alternative PAS (*C* allele); M4.*TP53*^A^: same as *TP53*^A^ except that 4 nucleotides were mutated to eliminate the miR-382 binding site; M4.*TP53*^C^: same as *TP53*^C^ except that 4 nucleotides were mutated to eliminate the miR-382 binding site; M4′.*TP53*^A^: same as *TP53*^A^ except that 4 nucleotides were mutated to eliminate the miR-325 binding site; M4′.*TP53*^C^: same as *TP53*^C^ except that 4 nucleotides were mutated to eliminate the miR-325 binding site; M8.*TP53*^A^: same as *TP53*^A^ except that 8 nucleotides were mutated to eliminate both binding sites; M8.*TP53*^C^: same as *TP53*^C^ except that 8 nucleotides were mutated to eliminate both binding sites. Biotinylated RNA fragments were transferred into *TP53*-null H1299 cells. Twenty four hours after transfection, cells were lysed followed by Pierce™ Streptavidin Magnetic Beads incubation and pull-down. After DNase I treatment, RNA purification, and reverse transcription, the relative amount of co-purified miR-125b (**b**), miR-325 (**c**) and miR-382 (**d**) were determined by qPCR. Values were normalized to the sample with the highest amount of miRNAs. *n* = 3 biological triplicates and *n* = 4-6 technical triplicates. **, *P* < 0.01; ***, *P* < 0.001. (**e**) Schematic diagram showing the synthesized 3′end biotin-labeled miRNAs and the 4 constructs expressing EGFP with the *TP53* 3′UTR (**Fig. S9**). **(f,g)** H1299 cells were mock transfected (Control), transfected with a luciferase report plasmid with *SV40* 3′UTR (*SV40*), transfected one of the EGFP expression plasmid, or co-transfected with one of the EGFP expression plasmid and one of the biotinylated miRNAs (**e**). Cells were lysed 24 h after transfection, Pierce™ Streptavidin Magnetic Beads were used to purify miRNA-interacting partners, and *TP53* 3′UTR levels were determined by qPCR. Values were normalized to that of miR-125b in cells with the *TP53*^A^ construct. *n* = 3 biological triplicates and *n* = 3-6 technical triplicates. ** *P* < 0.01.

**Fig. S11.**
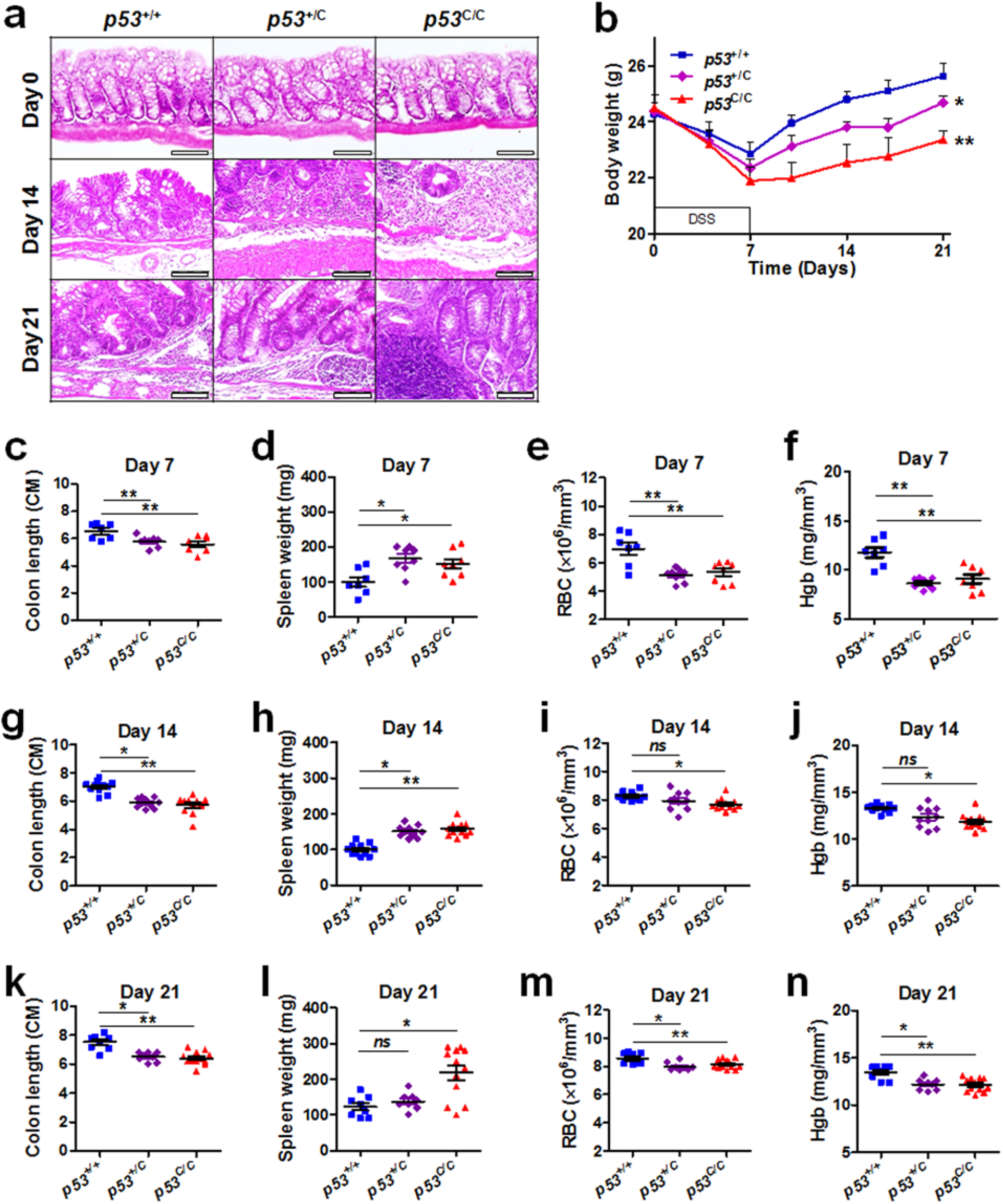
The *p53*^C^ allele promotes DSS-induced colitis in mice. Mice were treated with 2% dextran sulfate sodium (DSS) in the drinking water for 7 days and then given regular water; tissues were obtained at day 0 (no DSS treatment), day 7 (right after DSS treatment), day 14, and day 21. (**a**) Representative images for H&E staining of colon tissues. *n* ≥ 5 mice for each genotype. Scale bar, 100 µm. (**b**) Change in body weight of mice with 2% DSS treatment. Data were presented as mean ± s.e.m; *n* ≥ 10 mice for each genotype. * *P* < 0.05, ** *P* < 0.01. (**c-n**) Systemic inflammation and tissue damage in mice after DSS treatment. (**c,g,k**) Colon length, (**d,h,l**) Spleen weight, (**e,i,m**) Red blood cell (RBC), and (**f,j,n**) Hemoglobin (Hgb) in WT and mutant mice at day 7, 14, and 21. Data were presented as mean ± s.e.m; *n* ≥ 8 mice for each genotype. * *P* < 0.05, ** *P* < 0.01; *ns*, not significant.

**Fig. S12.**
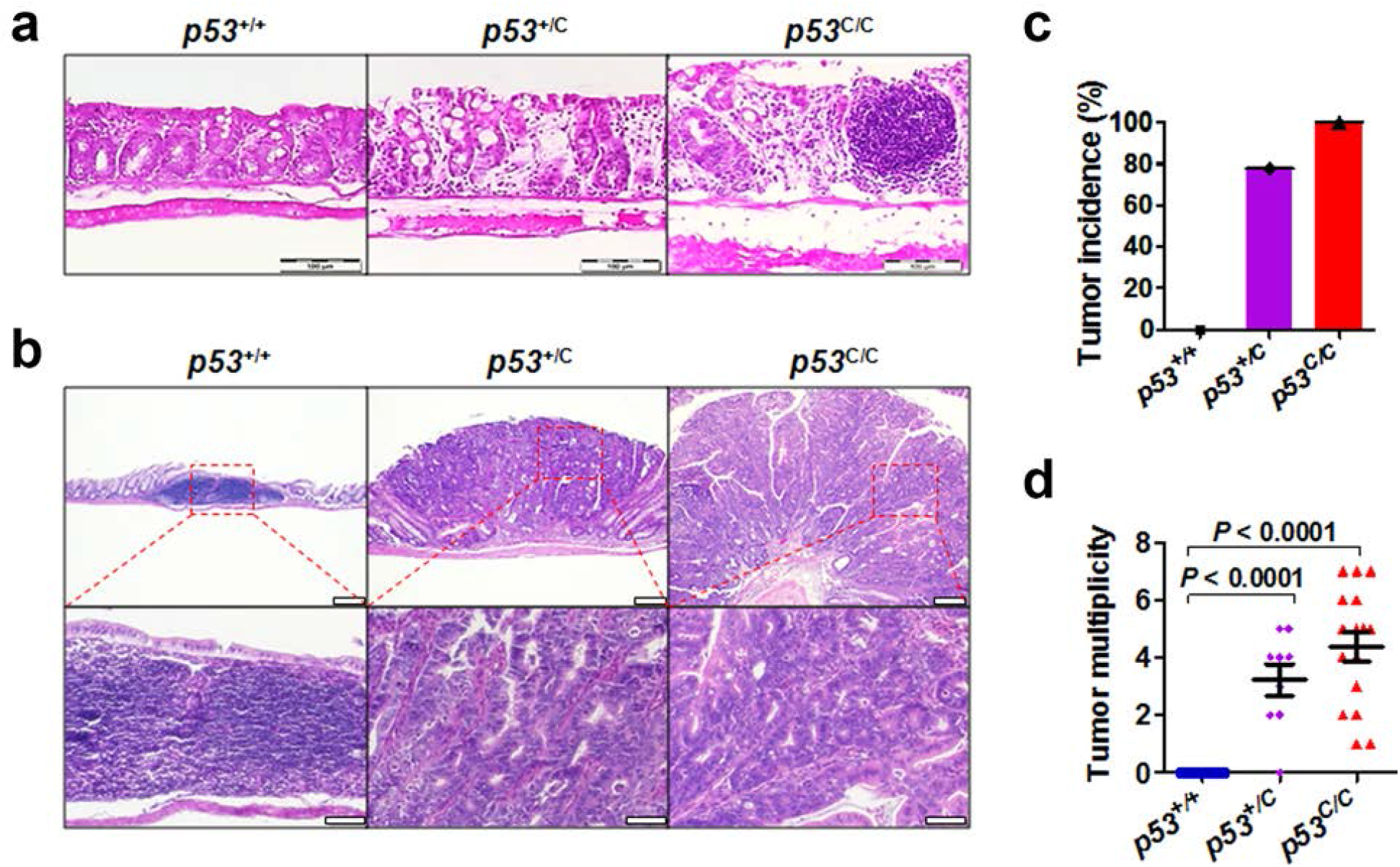
The *p53*^C^ allele accelerates colon carcinogenesis in mice. Mice were treated with a single dose of azoxymethane (AOM, 10 mg/kg, day 0) by intraperitoneal injection, followed by treatment with 2% DSS in drinking water for 7 days. (**a**) H&E staining of colon tissue sections from mice at day 7 after 2% DSS treatment only. Colon tissues were collected 8 weeks after DSS treatment. Scale bar, 100 µm. *n* ≥ 3 mice for each genotype. (**b**) Pathologic analyses of colon tissue lesions and colorectal adenomas. Colon tissues were collected 8 weeks after AOM-DSS treatment. Scale bar, 500 µm (top row), 100 µm (bottom row). *n* ≥ 3 mice for each genotype. (**c**) Tumor incidence for mice treated with AOM/DSS. *n* = 10, 9, and 16 for *p53*^+/+^, *p53*^+/C^, and *p53*^C/C^, respectively. (**d**) Tumor multiplicity in mice treated with AOM/DSS. Data were presented as mean ± s.e.m; *P* < 0.0001 (Mann-Whitney *U* test).

**Fig. S13.**
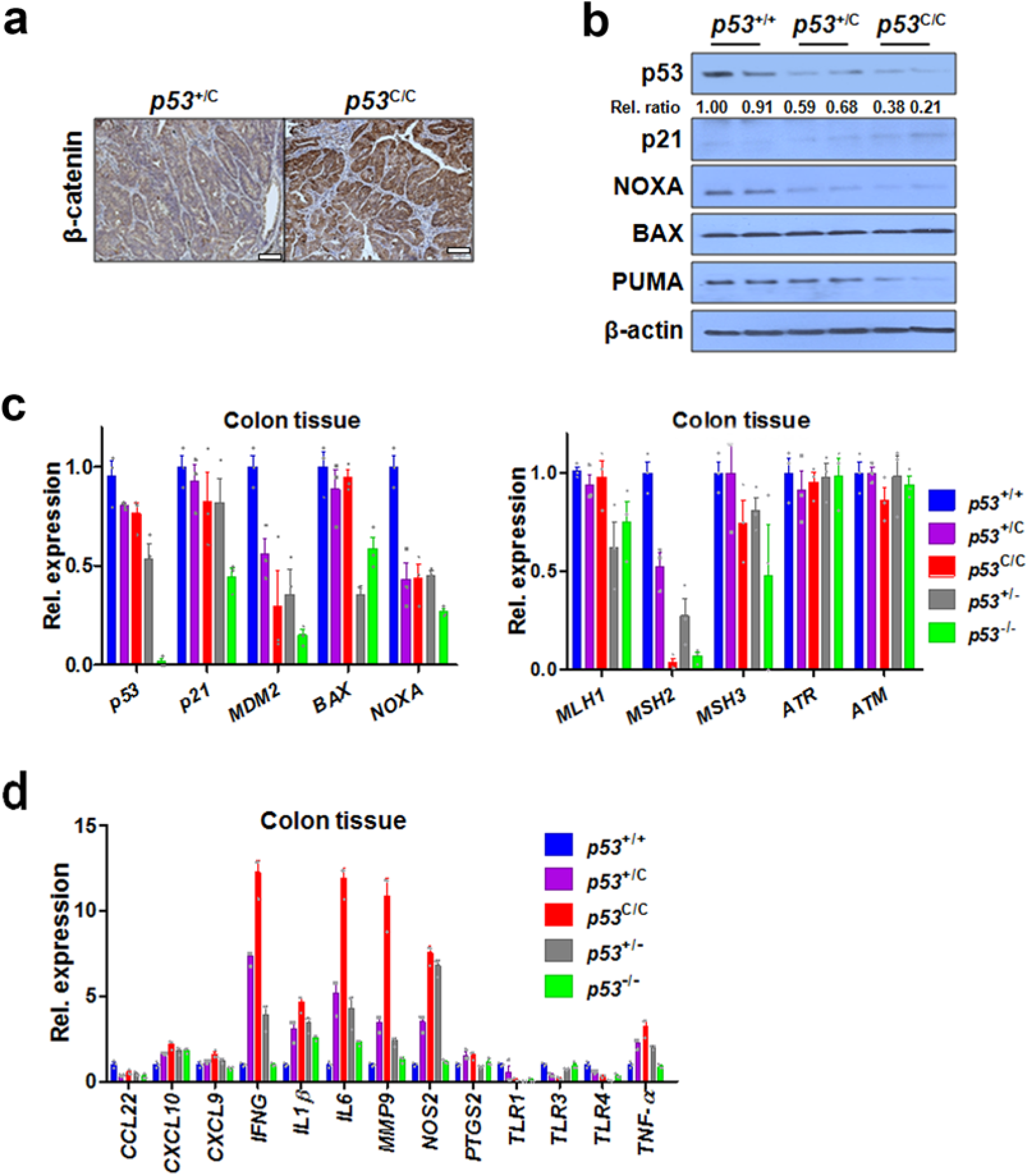
The *p53*^C^ allele compromises the p53 pathway in colon tissues. (**a**) IHC staining of β-catenin in colon tumors from mice treated with AOM/DSS. *n* = 3 mice for each genotype. Scale bar, 50 µm. (**b**) Western blotting analyses of the p53 pathway in mouse colon tissues. (c) qPCR assays for the expression of p53 pathway and DNA damage pathway genes in colon tissues from untreated mice with various *p53* alleles. Data were presented as mean ± s.e.m; *n* ≥ 3 mice for each genotype. (**d**) qPCR assay to determine the expression of inflammation genes in colon tissue from untreated mice with various *p53* alleles. Data were presented as mean ± s.e.m; *n* ≥ 3 mice for each genotype.

**Fig. S14.**
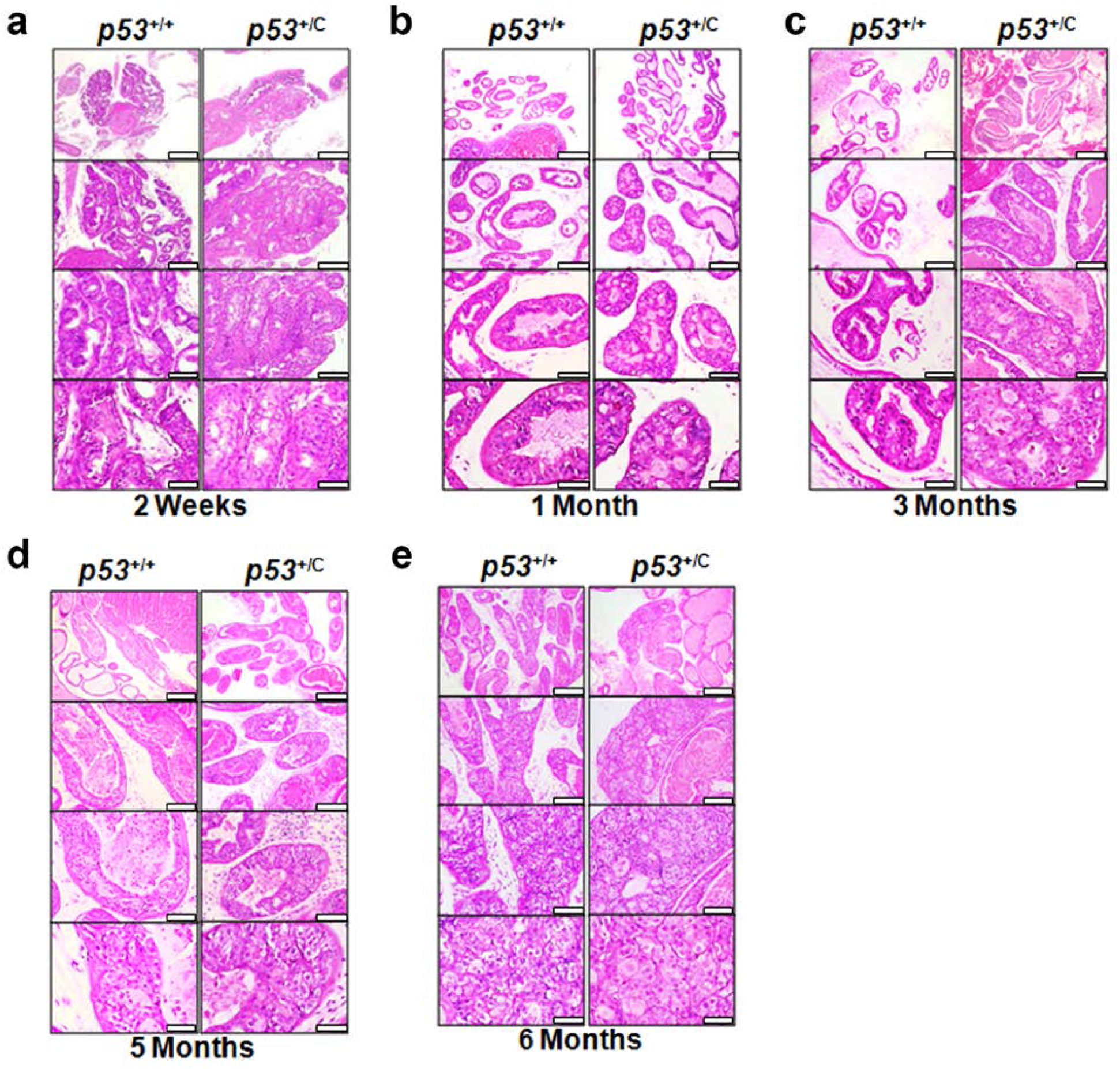
The *p53*^C^ allele accelerates prostate tumorigenesis in mice. F1 *Myc*-overexpressing male offspring of *p53*^+/C^ mice crossed with *Myc*-overexpressing mice (Hi-*Myc*, FVB- Tg(ARR2/Pbsn-MYC)7Key/ Nci) were used. (**a-e**) H&E staining of prostate tissue sections from mice at age of 2 weeks (**a**), 1 month (**b**), 3 months (**c**), 5 months (**d**), and 6 months (**e**). Scale bars: 200 µm (top row), 100 µm (second row), 50 µm (third row), and 20 µm (bottom row). *n* ≥3 mice for each genotype.

**Table S1.** rs78378222[C] and cancer risk.

| Cancers | MAF<sup>1</sup><br>(%) | Cases<br>(N) | Controls<br>(N) | OR<sup>2</sup> | P Value | References |
| --- | --- | --- | --- | --- | --- | --- |
| Glioma | 1.3 | 4,147 | 7,435 | 3.74 | $6.86 \times 10^{-24}$ | 13 |
| | 0.87-1.92 | 1,395 | 45,937 | 2.35 | $1.0 \times 10^{-5}$ | 8 |
| | 1.1 | 566 | 603 | 3.54 | $1.0 \times 10^{-4}$ | 12 |
| | 1.25 | 1,856 | 4,955 | 3.14 | $6.48 \times 10^{-11}$ | 14 |
| | 1.3 | 4,572 | 3,286 | 2.53 | $8.64 \times 10^{-38}$ | 17 |
| All brain tumors | 1.92 | 327 | 20,824 | 2.18 | $1.8 \times 10^{-3}$ | 8 |
| Neuroblastoma | 0.1-1.3 | 2,801 | 7,446 | 2.3 | $2.03 \times 10^{-11}$ | 18 |
| Esophageal squamous cell carcinoma | 1.0 | 405 | 810 | 3.22 | $1.34 \times 10^{-4}$ | 19 |
| Skin basal cell carcinoma | 0.46-1.92 | 4,319 | 51,810 | 2.16 | $2.2 \times 10^{-20}$ | 8 |
| Prostate cancer | 0.28-1.92 | 7,790 | 50,629 | 1.44 | $2.4 \times 10^{-6}$ | 8 |
| Colorectal adenoma | 1.85-1.92 | 4,095 | 43,022 | 1.39 | $1.6 \times 10^{-4}$ | 8 |
| Breast cancer | 0.5-1.91 | 4,985 | 42,995 | 1.06 | 0.57 | 8 |
| High-risk breast cancer<sup>3</sup> | 0.5-1.91 | 1,739 | 40,250 | 0.87 | 0.36 | 8 |
| Melanoma | 0.5-1.91 | 2,520 | 46,644 | 1.07 | 0.64 | 8 |
|  | 1.39 | 1,329 | 3,000 | 1.15 | 0.680 | 63 |
| Lung cancer | 1.04 | 1,013 | 3,000 | 0.84 | 0.379 | 63 |
| Squamous cell carcinoma of head and neck | 0.59 | 1,096 | 3,000 | 0.44 | 0.008 | 63 |
| Uterine leiomyoma | 1.8 | 16,595 | 523,330 | 1.74 | $4.0 \times 10^{-37}$ | 20 |
| All cancers (meta-analysis) |  | 36,599 | 91,272 | 1.51 | <0.001 | 64 |

<sup>1</sup> MAF, minor allele frequency in the control groups.
<sup>2</sup> OR, odds ratio.
<sup>3</sup> High-risk breast cancer is defined as breast cancer diagnosed before age of 50 or that having a history of multiple primary breast tumors.

**Table S2.**
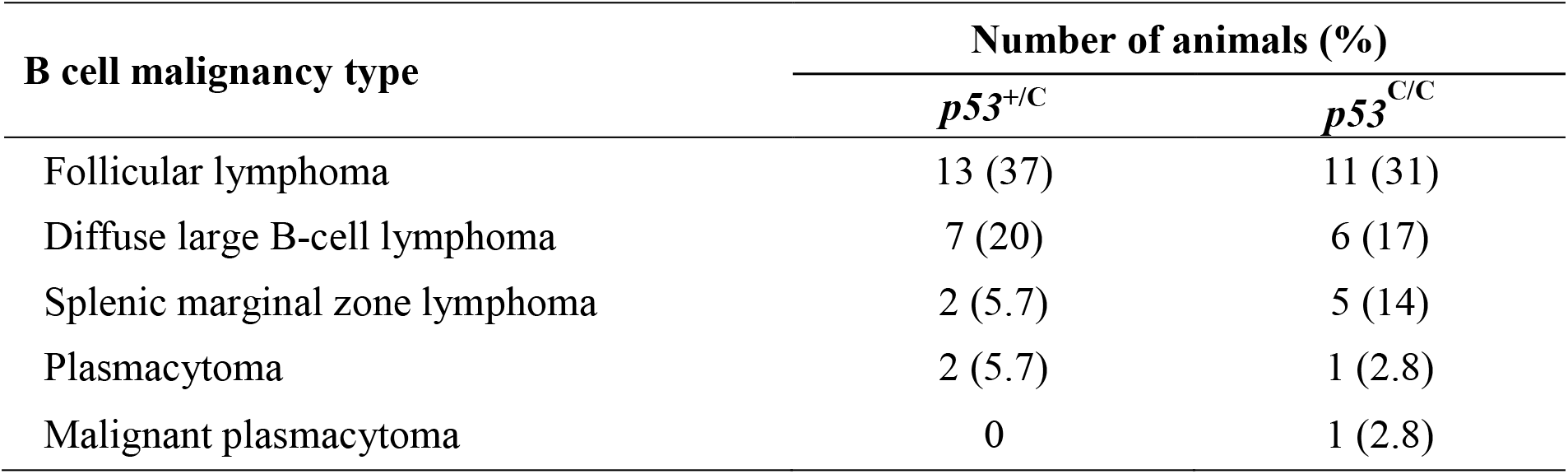
Summary of B-cell malignancies developed in mutant mice.

**Table S3.**
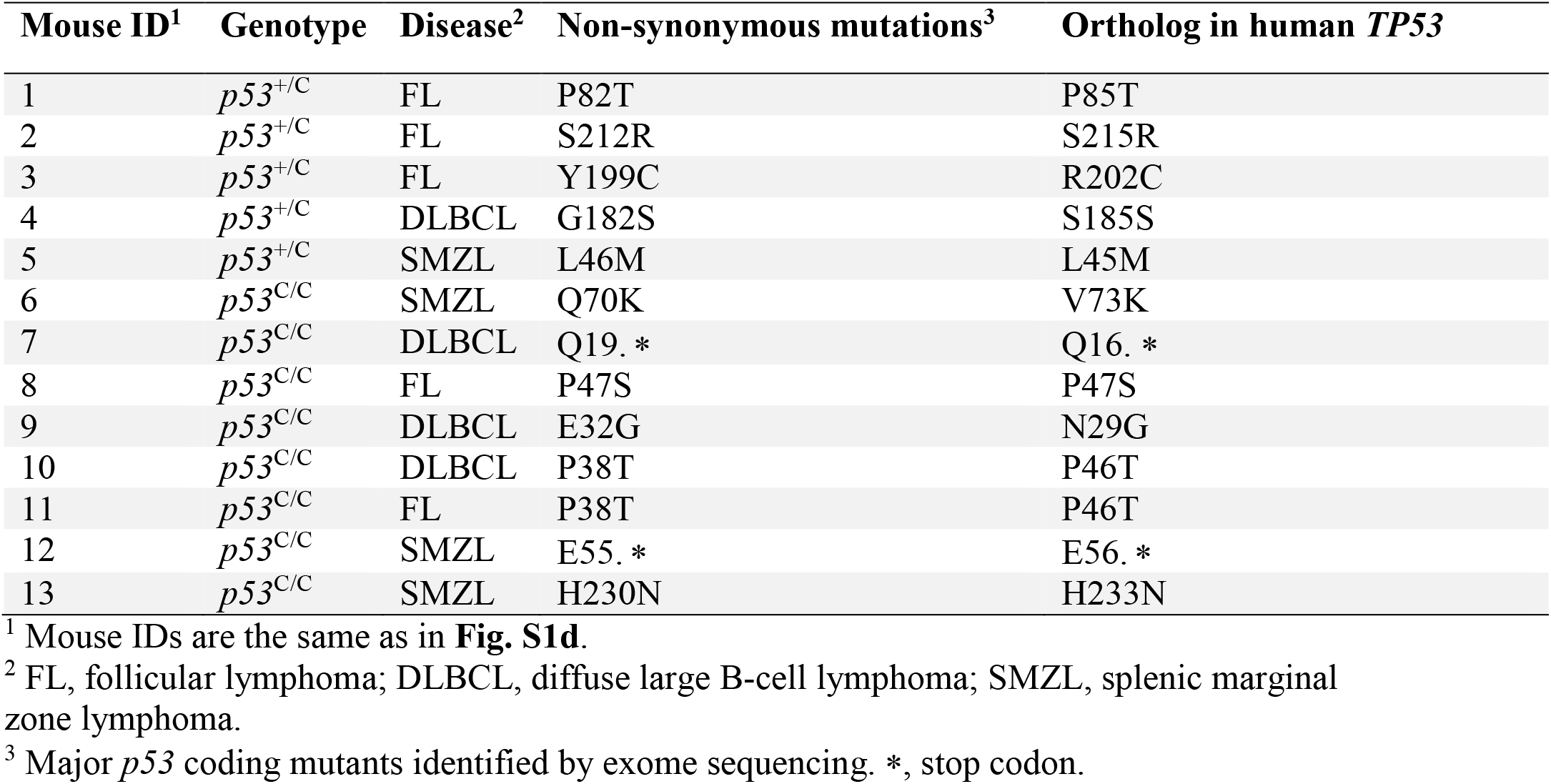
p53 coding mutations in tumors developed in p53^+/C^ and p53^C/C^ mice.

**Table S4.** Summary of 3-MCA-induced sarcoma development in mice.

| **3-MCA**<br>**dose (μg)** | ***p53***<br>**Genotype** | **Number**<br>**of mice** | **Sarcoma**<br>**(%)** | **Earliest**<br>**tumor**<br>**(days)** | **Median time to**<br>**tumor development**<br>**(days)** | **Follow-**<br>**up (days)** |
| --- | --- | --- | --- | --- | --- | --- |
| 50 | *p53*<sup>+/+</sup> | 9 | 78 | 115 | 130 | 200 |
|  | *p53*<sup>+/*C*</sup> | 6 | 100 | 108 | 117.5 | 135 |
|  | *p53*<sup>*C/C*</sup> | 8 | 100 | 80 | 114 | 128 |
| 300 | *p53*<sup>+/+</sup> | 9 | 100 | 94 | 115 | 122 |
|  | *p53*<sup>+/*C*</sup> | 6 | 100 | 87 | 104 | 115 |
|  | *p53*<sup>*C/C*</sup> | 10 | 100 | 66 | 96.5 | 115 |

**Table S5.** Mouse line crosses and genotypes selected.

| Parent genotype<br>(Genetic background) | Offspring genotype | Offspring genetic<br>background | Offspring<br>Selected<br>for study |
| --- | --- | --- | --- |
| $p53^{+/C} \times p53^{+/C}$<br>(C57 × C57) | $p53^{+/+}$ | C57 | Yes |
| | $p53^{+/C}$ | C57 | Yes |
| | $p53^{C/C}$ | C57 | Yes |
| $p53^{+/-} \times p53^{+/-}$<br>(C57 × C57) | $p53^{+/+}$ | C57 | Yes |
| | $p53^{+/-}$ | C57 | Yes |
| | $p53^{-/-}$ | C57 | Yes |
| $p53^{+/C} \times PyVT *$<br>(C57 × FVB) | $p53^{+/+}; PyVT$ | 50% FVB +50% C57 | Yes |
| | $p53^{+/C}; PyVT$ | 50% FVB +50% C57 | Yes |
| | $p53^{+/+}$ | 50% FVB +50% C57 | No |
| | $p53^{+/C}$ | 50% FVB +50% C57 | No |
| $p53^{+/C} \times p53^{+/C}; PyVT *$<br>(C57 × [50% FVB +50% C57]) | $p53^{+/+}; PyVT$ | 25% FVB +75% C57 | Yes |
| | $p53^{+/C}; PyVT$ | 25% FVB +75% C57 | Yes |
| | $p53^{C/C}; PyVT$ | 25% FVB +75% C57 | Yes |
| | $p53^{+/+}$ | 25% FVB +75% C57 | No |
| | $p53^{+/C}$ | 25% FVB +75% C57 | No |
| | $p53^{C/C}$ | 25% FVB +75% C57 | No |
| $p53^{+/C} \times Erbb2 *$<br>(C57 × FVB) | $p53^{+/+}; Erbb2$ | 50% FVB +50% C57 | Yes |
| | $p53^{+/C}; Erbb2$ | 50% FVB +50% C57 | Yes |
| | $p53^{+/+}$ | 50% FVB +50% C57 | No |
| | $p53^{+/C}$ | 50% FVB +50% C57 | No |
| $p53^{+/-} \times Erbb2 *$<br>(C57 × FVB) | $p53^{+/+}; Erbb2$ | 50% FVB +50% C57 | Yes |
| | $p53^{+/-}; Erbb2$ | 50% FVB +50% C57 | Yes |
| | $p53^{+/+}$ | 50% FVB +50% C57 | No |
| | $p53^{+/-}$ | 50% FVB +50% C57 | No |
| $p53^{+/C} \times Hi-Myc *$<br>(C57 × FVB) | $p53^{+/+}; Hi-Myc$ | 50% FVB +50% C57 | Yes |
| | $p53^{+/C}; Hi-Myc$ | 50% FVB +50% C57 | Yes |
| | $p53^{+/+}$ | 50% FVB +50% C57 | No |
| | $p53^{+/C}$ | 50% FVB +50% C57 | No |

\* Offspring of these crosses may or may not carry the indicated transgene (*PyVT*, *Erbb2*, or *Hi-Myc*). C57, C57bl/6.

